# Protein interaction energy landscapes are shaped by functional and also non-functional partners

**DOI:** 10.1101/298174

**Authors:** Hugo Schweke, Marie-Hélène Mucchielli, Sophie Sacquin-Mora, Wanying Bei, Anne Lopes

## Abstract

In the crowded cell, a strong selective pressure operates on the proteome to limit the competition between functional and non-functional protein-protein interactions. We developed an original theoretical framework in order to interrogate how this competition constrains the behavior of proteins with respect to their partners or random encounters. Our theoretical framework relies on a two-dimensional (2D) representation of interaction energy landscapes with 2D energy maps that reflect in a synthetic way the propensity of a protein to interact with another protein. We investigated the propensity of protein surfaces to interact with functional and arbitrary partners and asked whether their interaction propensity is conserved during the evolution. Therefore, we performed several thousands of cross-docking simulations to systematically characterize the whole energy landscapes of 74 proteins interacting with different sets of homologs, corresponding to their functional partner’s family or arbitrary protein families. Then, we systematically compared the energy maps resulting from the docking of a given protein with the different protein families of the dataset. Strikingly, we show that the interaction propensity not only of the binding site but also of the rest of the protein surface is conserved for docking partners belonging to the same protein family. Interestingly, this observation holds for docked proteins corresponding to true but also to arbitrary partners. Our theoretical framework enables the characterization of the energy behavior of a protein in interaction with hundreds of selected partners and opens the way for further developments to study the behavior of proteins in a specific environment.

## Introduction

Biomolecular interactions are central for many physiological processes and are of utmost importance for the functioning of the cell. Particularly protein-protein interactions have attracted a wealth of studies these last decades [1–5]. The concentration of proteins in a cell has been estimated to be approximately 2-4 million proteins per cubic micron [6]. In such a highly crowded environment, proteins constantly encounter each other and numerous non-specific interactions are likely to occur [7–10]. For example, in the cytosol of *S. cerevisiae* a protein can encounter up to 2000 different proteins [11]. In this complex jigsaw puzzle, each protein has evolved to bind the right piece(s) in the right way (positive design) and to prevent misassembly and non-functional interactions (negative design) [12–16]).

Consequently, positive design constrains the physico-chemical properties and the evolution of protein-protein interfaces. Indeed, a strong selection pressure operates on binding sites to maintain the functional assembly including the functional partner and the functional binding mode. For example, homologs sharing at least 30% sequence identity almost invariably interact in the same way [17]. Conversely, negative design prevents proteins to be trapped in the numerous competing non-functional interactions inherent to the crowded environment of the cell. Many studies were reported on the relationship between the propensity of proteins for promiscuous interactions and their abundances or surface properties [18–21]. Particularly, it has been shown that the misinteraction avoidance shapes the evolution and physico-chemical properties of abundant proteins, resulting in a slower evolution and less sticky surfaces than what is observed for less abundant ones [18,22–26]. The whole surface of abundant proteins is thus constrained, preventing them to engage deleterious non-specific interactions that could be of dramatic impact for the cell at high concentration [25]. Recently, it has been shown in *E. coli* that the net charge as well as the charge distribution on protein surfaces affect the diffusion coefficients of proteins in the cytoplasm [19,27]. Positively charged proteins move up to 100 times more slowly as they get caught in non-specific interactions with ribosomes which are negatively charged and therefore, shape the composition of the cytoplasmic proteome [27].

All these studies show that both positive and negative design effectively operate on the whole protein surface. Binding sites are constrained to maintain functional assemblies (i.e. functional binding modes and functional partners) while the rest of the surface is constrained to avoid non-functional assemblies. Consequently, these constraints should shape the energy landscapes of functional but also non-functional interactions so that non-functional interactions do not prevail over functional ones. This should have consequences (i) on the evolution of the propensity of a protein to interact with its environment (including functional and non-functional partners) and (ii) on the evolution of the interaction propensity of the whole surface of proteins, non-interacting surfaces being in constant competition with functional binding sites. We can hypothesize that the interaction propensity of the whole surface of proteins is constrained during evolution in order to (i) ensure that proteins correctly bind functional partners, and (ii) limit non-functional assemblies as well as interactions with non-functional partners.

In this work, we focus on protein surfaces as a proxy for functional and non-functional protein-protein interactions. We investigate their interaction energy landscapes with native and non-native partners and ask whether their interaction propensity is conserved during evolution. With this aim in mind, we performed large-scale docking simulations to characterize interactions involving either native or native-related (i.e. partners of their homologs) partners or arbitrary partners. Docking simulations enable the characterization of all possible interactions involving either functional or arbitrary partners, and thus to simulate the interaction of arbitrary partners which is very difficult to address with experimental approaches. Docking algorithms are now fast enough for large-scale applications and allow for the characterization of interaction energy landscapes for thousand of protein couples. Typically, a docking simulation takes from a few minutes to a couple of hours on modern processors [28–30], opening the way for extensive cross-docking experiments [31–35]. Protein docking enables the exploration of the interaction propensity of the whole protein surface by simulating alternative binding modes. Here, we performed a cross-docking experiment involving 74 selected proteins docked with their native-related partners and their corresponding homologs, as well as arbitrary partners and their corresponding homologs. We represented the interaction energy landscapes resulting from each docking calculation with a two dimensional (2D) energy map in order to (i) characterize the propensity of all surface regions of a protein to interact with a given partner (either native-related or not) and (ii) easily compare the energy maps resulting from the docking of a same protein with different sets of homologous partners, thus addressing the evolution of the propensity of a protein to interact with homologous partners either native or arbitrary.

## Results

### The interaction propensity of the whole surface of the human ubiquitin carboxyl-terminal hydrolase 14 is conserved for homologous protein ligands, be they functional partners or random encounters

If positive and negative design constraint the propensity of the whole surface of proteins to interact with their functional partners or random encounters, this should shape the evolution of interaction energy landscapes of functional protein pairs but also of random encounter pairs. Consequently, we expect that the interaction energy landscape involving a protein pair (functional or arbitrary) is conserved for a homologous pair. Testing this hypothesis involves being able to characterize the interaction propensity of the whole surface of a protein. Therefore we designed a procedure based on a two-dimensional (2D) representation of docking energy landscapes with 2D energy maps which reflect the propensity of a protein (i.e. the receptor) to interact with the docked partner (i.e. the ligand) (*Materials and Methods*, Fig 1A-C). The procedure is asymmetrical and the resulting energy map provides the distribution of all docking energies over the whole receptor surface thus reflecting the propensity of the receptor to interact with the docked ligand. Fig 2 represents the energy maps computed for the receptor 2AYN_A, the human ubiquitin carboxyl-terminal hydrolase 14 (family UCH) docked with (i) its native partner (1XD3_B, ubiquitin-related family), a homolog of its partner (defined as a native-related partner) (1NDD_B) and (ii) two arbitrary homologous ligands (1YVB_A and 1NQD_B from the papain-like family). For all four ligands, either native-related or arbitrary partners, docking calculations lead to an accumulation of low-energy solutions (hot regions in red) around the two experimentally known binding sites of the receptor. The first one corresponds to the interaction site with the native partner, ubiquitin (pdb id 2ayo). The second one corresponds to its homodimerisation site (pdb id 2ayn). This indicates that native-related but also arbitrary partners tend to bind onto the native binding sites of native partners as observed in earlier studies [34,36]. Indeed, the low energy solutions tend to accumulate systematically in the vicinity of the two native interaction sites. Whereas the low energy solutions obtained for both ligand families accumulate around the native binding sites of 2AYN_A, the two ligand families display clear differences in the rest of the map. Indeed, the energy maps obtained with the ligands of the ubiquitin-like family both reveal two sharp hot regions around the native sites and a subset of well-defined cold regions (i.e. blue regions corresponding to high energy solutions) placed in the same area in the map’s upper-right quadrant. In contrast, the energy maps obtained for the ligands of the papain-like family display a large hot region around the two native binding sites of the receptor, extending to the upper-left and bottom-right regions of the map and suggesting a large promiscuous binding region for these ligands. The interaction propensity of the two binding sites of 2AYN_A but also of the other regions of its surface seems to be conserved for homologous ligands and specific to each ligand family whether the ligands correspond to native-related partners or not (Fig 2).

**Fig. 1.**
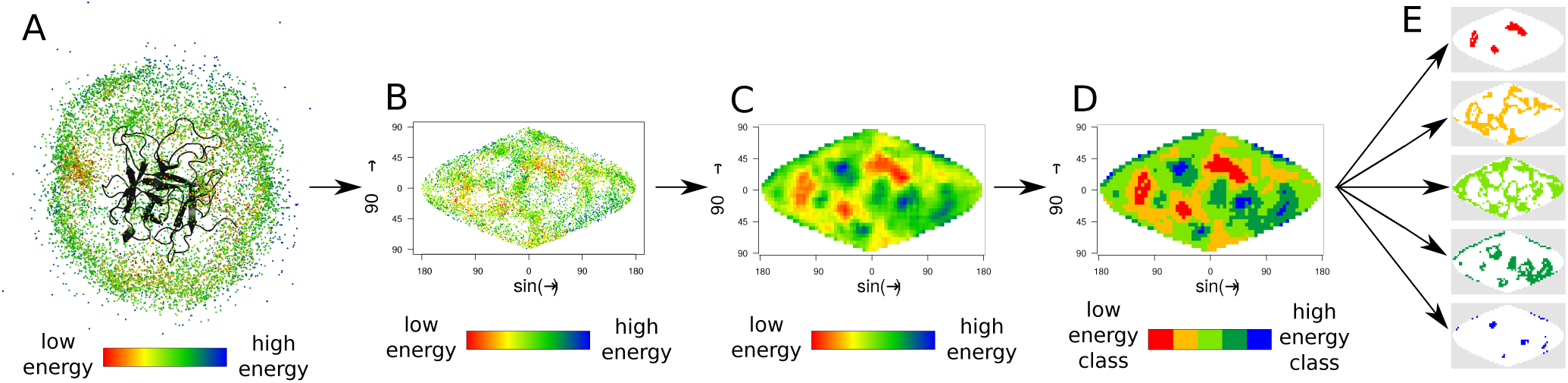
2D asymmetrical representation of docking energy landscapes and resulting energy maps. (*A*) Three-dimensional (3D) representation of the ligand docking poses around the receptor. Each dot corresponds to the center of mass (CM) of a ligand docking pose and is colored according to its docking energy score. (*B*) Representation of the CM of the ligand docking poses after an equal-area 2D sinusoidal projection. CMs are colored according to the same scale as in A. (*C*) Continuous energy map (see *Materials and Methods* for more details). (*D*) Five-class map. The energy map is discretized into five energy classes (*E*) One-class maps. Top to bottom: one-class maps that highlight respectively hot, warm, lukewarm, cool and cold regions.

**Fig. 2.**
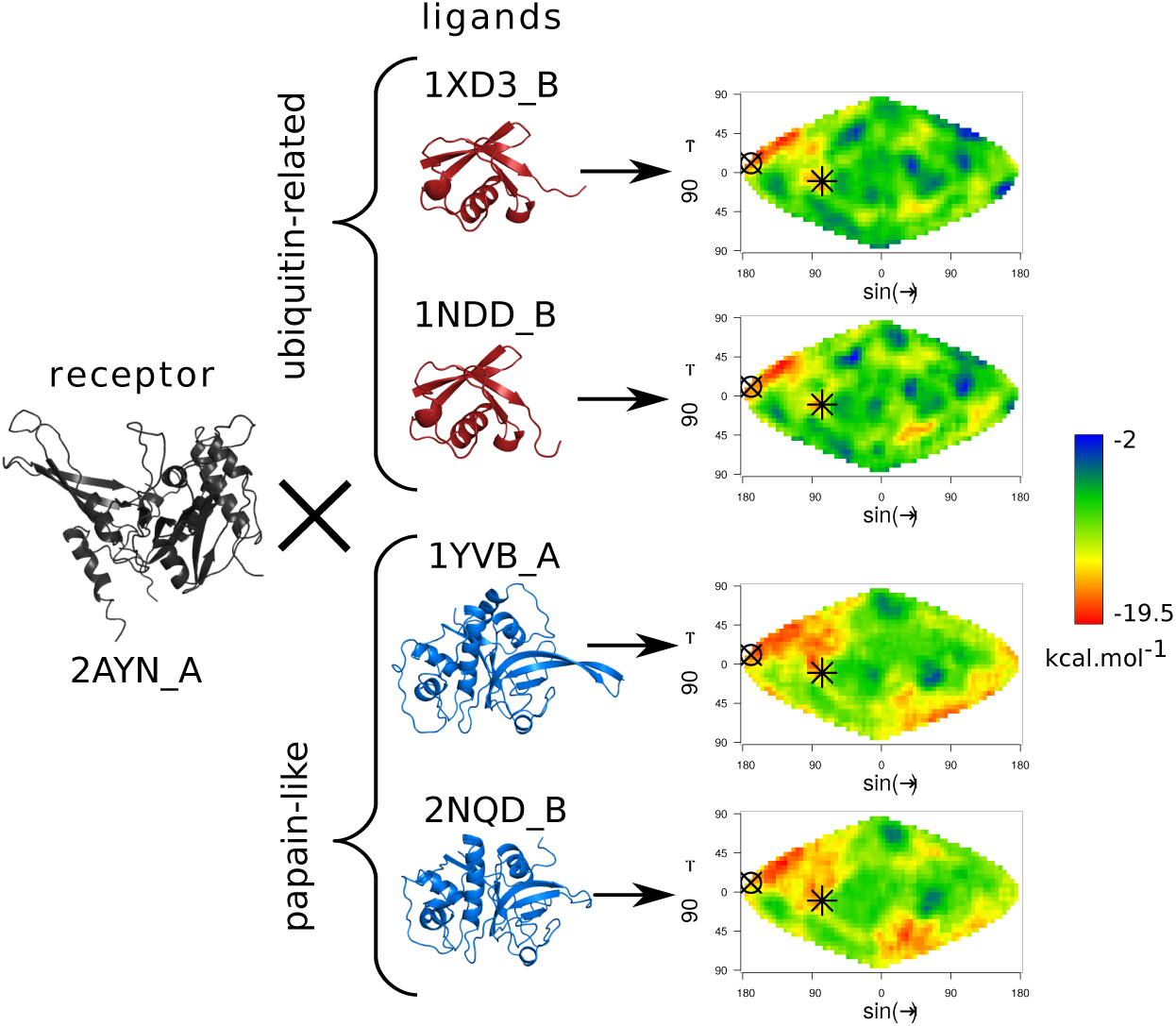
Interaction propensity for the receptor 2AYN_A and four different ligands. 2D energy maps for the receptor 2AYN_A (ubiquitin carboxyl-terminal hydrolase (UCH) family) docked with the ligands 1XD3_B (native partner), 1NDD_B (homolog of the native partner), 1YVB_A and 2NQD_B (arbitrary partners). The star indicates the localization of the experimentally determined interaction site for the ubiquitin, the circle-cross indicates the homodimerization site of 2AYN_A.

### Generalization to a large set of proteins

We asked whether this observation could be generalized to a large set of proteins. Therefore we built a database comprising 74 protein structures divided into 12 families of homologs (S1 Table and *Materials and Methods*). Each family displays different degrees of structural variability and sequence divergence in order to see the impact of these properties on the conservation of the interaction propensity inside a protein family. Each family has at least one native-related partner family (S1 Fig). For a protein A, we refer as native-related partners its native partner (when its three dimensional (3D) structure is available) and native partners of proteins that are homologous to the protein A. Arbitrary pairs refer to pairs of proteins for which no interaction has been experimentally characterized in the Protein Data Bank neither for their respective homologs [37]. Docking calculations are performed with the ATTRACT software [30]. Each protein (namely the receptor) is docked with the 74 proteins (namely the ligands) of the dataset (Fig 3A and *Materials and Methods*) and the 74 corresponding energy maps are calculated (Fig 3B and *Materials and Methods*). The 74 resulting energy maps are compared together with a Manhattan distance and all the energy map distances are stored in an energy map distance (EMD) matrix (Fig 3C and *Materials and Methods*). Each matrix entry (*i,j*) corresponds to the distance d_i,j_ between the energy maps of ligands *i* and *j* docked with a receptor *k* (Fig 3C and *Materials and Methods*). Consequently, a small distance d_i,j_ between ligands *i* and *j* docked with a receptor *k*, reflects a high similarity of their energy maps. In other words, the interaction propensity of the surface of the receptor *k* is similar for both ligands *i* and *j*. One should notice that energy maps computed for two unrelated receptors are not comparable since their surfaces are not comparable as well. Therefore, the procedure is asymmetrical and receptor-centered. It only compares energy maps calculated for different ligands docked with the same receptor. In order to prevent any bias from the choice of the receptor, each of the 74 proteins plays alternately the role of receptor and ligand. Consequently, the protocol presented in Fig 3 is repeated for the entire dataset where each protein plays the role of the receptor and is docked with the 74 proteins that play the role of ligands, thus resulting in 74 EMD matrices. In order to quantify the extent to which the interaction propensity of a receptor is conserved for homologous ligands, we evaluated to what extent distances calculated between homologous ligand pairs were smaller than distances calculated between random pairs. Fig 4 represents the boxplots of energy map distances calculated between random ligand pairs or between homologous ligand pairs docked with their native-related receptor or with the other receptors of the dataset. Homologous ligands display significantly lower energy map distances than random ligand pairs (Wilcoxon test *p* = 0) indicating that energy maps produced by homologous ligands are more similar than those produced by non-homologous ligands. Interestingly, this observation holds whether the receptor-ligand pair is a native pair or not. This suggests that the interaction propensity of a receptor is conserved for homologous partners be they native-related or not.

**Fig. 3.**
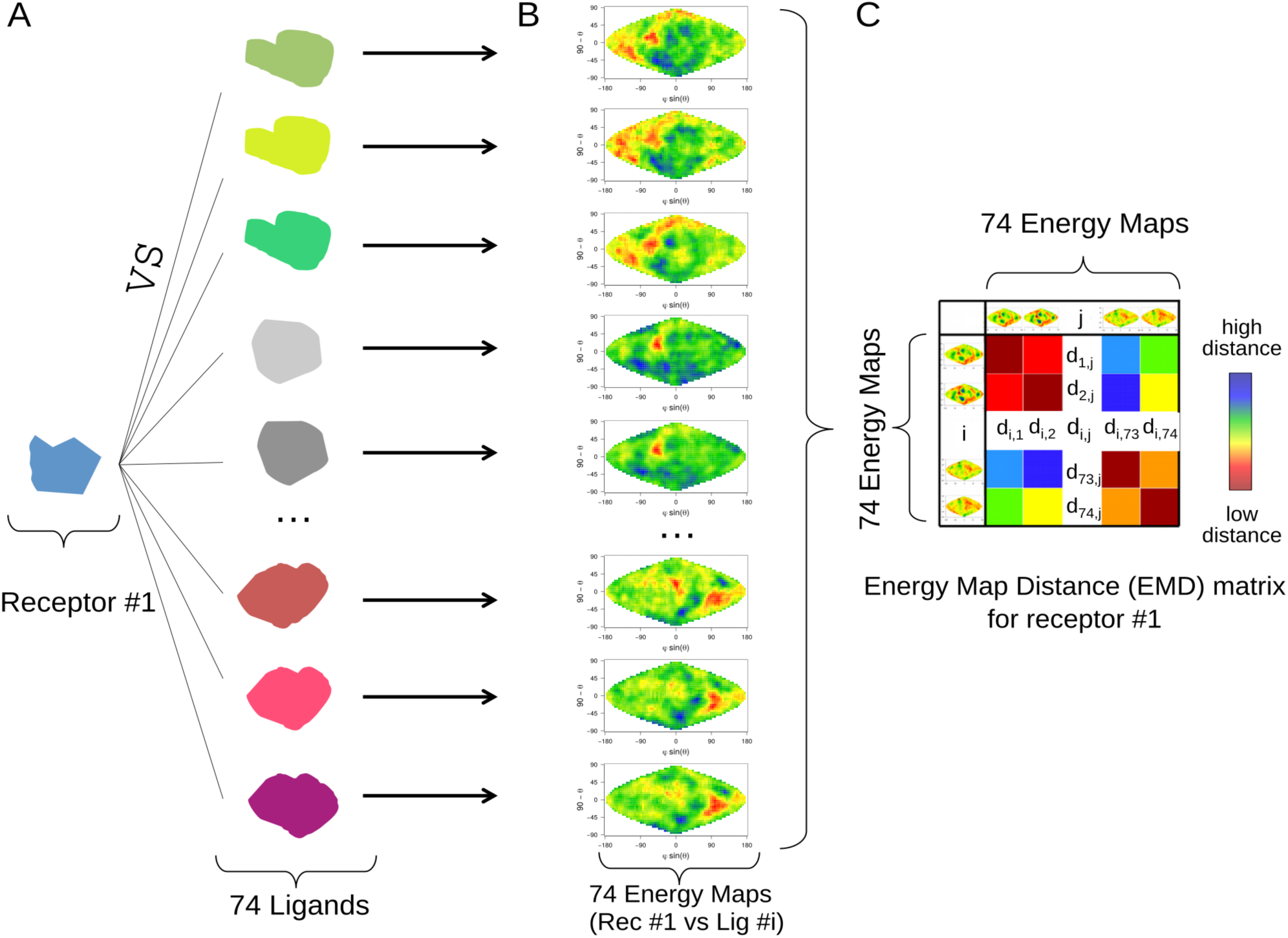
Experimental Protocol. (*A*) A receptor protein is docked with all proteins of the dataset (namely the ligands) resulting in 74 docking calculations. (*B*) For each docking calculation, an energy map is computed as well as its corresponding five-classes and one-class energy maps, with the procedure described in Fig 1 and *Materials and Methods*. (*C*) An energy map distance (EMD) matrix is computed, representing the pairwise distances between the 74 energy maps resulting from the docking of all ligands with this receptor. Each cell (*i,j*) of the matrix represents the Manhattan distance between the two energy maps resulting from the docking of ligands *i* and *j* with the receptor. A small distance indicates that the ligands *i* and *j* produce similar energy maps when docked with this receptor. In other words, it reflects that the interaction propensity of this receptor is similar for these two ligands. To prevent any bias from the choice of the receptor, the whole procedure is repeated for each receptor of the database, leading to 74 EMD matrices.

**Fig. 4.**
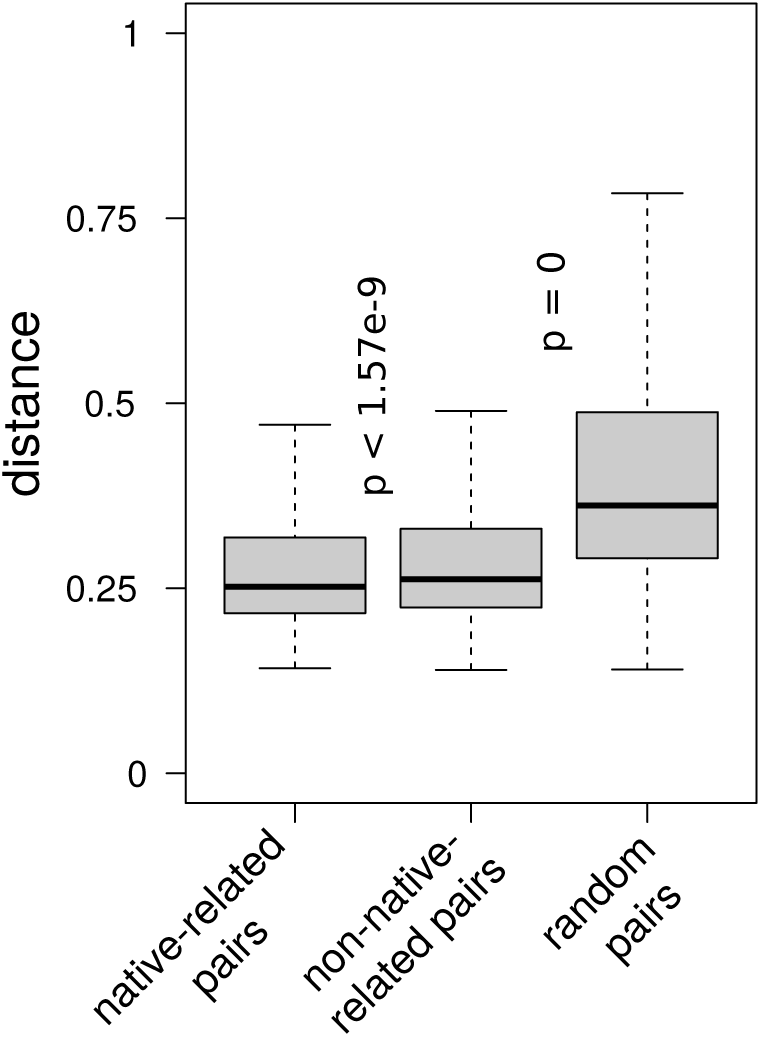
Boxplots of energy map pairwise distances between homologous ligand pairs from native-related partner families, homologous ligand pairs from arbitrary partner families and random ligand pairs. For each receptor, we computed (i) the average of energy map distances of pair of homologous ligands belonging to its native-related partner family(ies), (ii) the average of energy map distances of pair of homologous ligands belonging to its non-native-related partner families, and (iii) the average of energy map distances of random pairs. P-values are calculated with a unilateral Wilcoxon test.

### Energy maps are specific to protein families

The results presented above prompted us to assess the extent to which the interaction propensity of a receptor is specific to the ligand families it interacts with. If so, a receptor should lead to energy maps that are specific to the different ligand families and we should be able to retrieve homology relationships of ligands solely from the comparison of their energy maps. Therefore, we tested our ability to predict the homologs of a given ligand based only on the comparison of its energy maps with those of the other ligands. In order to prevent any bias from the choice of the receptor, the 74 EMD matrices are averaged in an averaged distances matrix (ADM) (see *Materials and Methods*). Each entry *(i,j)* of the ADM corresponds to the averaged distance between two sets of 74 energy maps produced by two ligands *i* and *j*. A low distance indicates that the two ligands display similar energy maps whatever the receptor is. We computed a receiver operating characteristic (ROC) curve from the ADM (see *Materials and Methods*) which evaluates our capacity to discriminate the homologs of a given ligand from non-homologous ligands by comparing their respective energy maps computed with all 74 receptors of the dataset. The true positive set consists in the homologous protein pairs while the true negative set consists in any homology-unrelated protein pair. The resulting Area Under the Curve (AUC) is equal to 0.79 (Fig 5). We evaluated the robustness of the ligand’s homologs prediction depending on the size of the receptor subset with a bootstrap procedure by randomly removing receptor subsets of different sizes (from 1 to 73 receptors). The resulting AUCs range from 0.77 to 0.79, and show that from a subset size of five receptors, the resulting prediction accuracy no longer significantly varies (risk of wrongly rejecting the equality of two variances (F-test) >5%), and is robust to the nature of the receptor subset (S2 Fig). Finally, we evaluated the robustness of the predictions according to the number of grid cells composing the energy maps. Therefore, we repeated the procedure using energy maps with resolutions ranging from 144×72 to 48×24 cells. S2 Table presents the AUCs calculated with different grid resolutions. The resulting AUCs range from 0.78 to 0.8 showing that the grid resolution has a weak influence on the map comparison. All together, these results indicate that homology relationships between protein ligands can be detected solely on the basis of the comparison of their energy maps. In other words, the energy maps calculated for a receptor docked with a set of ligands belonging to a same family are specific to this family. Interestingly, this observation holds for families displaying important sequence variations (S1 Table). For example, the AUC computed for the UCH and ubiquitin-related families are 0.98 and 0.88 respectively despite the fact that the average sequence identity of these families does not exceed 45% (S3 Fig and S1 Table). This indicates that energy maps are similar even for homologous ligands displaying large sequence variations.

**Fig. 5.**
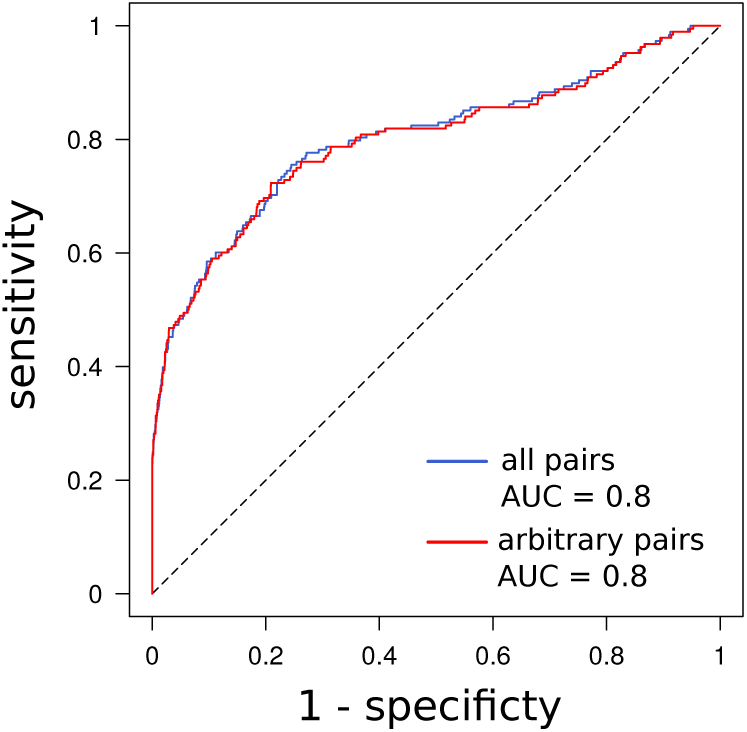
Receiver operating characteristic (ROC) curve and its Area Under the Curve (AUC). ROC are calculated on the averaged distance matrix (ADM) including either all pairs (blue) or only arbitrary pairs (red) (see *Materials and Methods* for more details).

We then specifically investigated the energy maps of each family in order to see whether some ligands behave energetically differently from their family members. On the 74 ligands, only five (2L7R_A, 4BNR_A, 1BZX_A, 1QA9_A, 1YAL_B) display energy maps that are significantly different from those of their related homologs (Z-tests *p-values* for the comparison of the averaged distance of each ligand with their homologs versus the averaged distance of all ligands with their homologous ligands ≤ 5%). In order to identify the factors leading to differences between energy maps involving homologous ligands, we computed the pairwise sequence identity and the root mean square deviation (RMSD) between the members of each family. Interestingly, none of these criteria can explain the energy map differences observed within families (Fisher test *p* of the linear model estimated on all protein families >0.1) (see Fig 6B-C for the ubiquitin-related family, S4-S14B-C Fig for the other families, and S3 Table for details). Fig 6A represents a subsection of the ADM for the ubiquitin-related family (i.e. the energy map distances computed between all the members of the ubiquitin-like family and averaged over the 74 receptors). Low distances reflect pairs of ligands with similar energy behaviors (i.e. producing similar energy maps when interacting with a same receptor) while high distances reveal pairs of ligands with different energy behaviors. 2L7R_A distinguishes itself from the rest of the family, displaying high-energy map distances with all of its homologs. RMSD and sequence identity contribute modestly to the energy map distances observed in Fig 6A (Spearman correlation test *p^RMSD^* = 0.01 and *p^seq^* = 0.02 (S3 Table, Fig 6B-C)). Fig 6D shows a projection of the electrostatic potential calculated with APBS [38] on the surface of the seven ubiquitin-related family members (for more details, see S15 Fig and *Materials and Methods*). Fig 6E represents the electrostatic maps distances computed between all members of the family. 2L7R_A clearly stands out, displaying a negative electrostatic potential over the whole surface while its homologs harbor a remarkable fifty-fifty electrostatic distribution (Fig 6D). The negatively charged surface of 2L7R_A is explained by the absence of the numerous lysines that are present in the others members of the family (referred by black stars, Fig 6D). Lysines are known to be essential for ubiquitin function, enabling the formation of polyubiquitin chains on target proteins. Among the seven lysines of the ubiquitin, K63 polyubiquitin chains are known to act in non-proteolytic events while K48, K11, and the four other lysines polyubiquitin chains are presumed to be involved into addressing proteins to the proteasome [39]. 2L7R_A is a soluble UBL domain resulting from the cleavage of the fusion protein FAU [40]. Its function is unrelated to proteasomal degradation, which might explain the lack of lysines on its surface and the differences observed in its energy maps. Interestingly, the differences observed for the energy maps of 1YAL_B (Papain-like family) (S4 Fig) and 4BNR_A (eukaryotic proteases family) (S5 Fig) regarding their related homologs can be explained by the fact that they both display a highly charged surface. These two proteins are thermostable [41,42], which is not the case for their related homologs, and probably explains the differences observed in their relative energy maps. The V-set domain family is split into two major subgroups according to their averaged energy map distances (S6A Fig). The first group corresponds to CD2 proteins (1QA9_A and its unbound form 1HNF_A) and differs significantly from the second group (Z-test *p =* 0.03 and *p* = 0.05 respectively). The second group corresponds to CD58 (1QA9_B and its unbound form 1CCZ_A) and CD48 proteins (2PTT_A). Interestingly, CD2 is known to interact with its homologs (namely CD58 and CD48) through an interface with a striking electrostatic complementarity [43]. The two subgroups have thus evolved distinct and specific binding sites to interact together. We can hypothesize that they have different interaction propensities resulting in the differences observed between their corresponding energy maps. These five cases illustrate the capacity of our theoretical framework to reveal functional or biophysical specificities of homologous proteins that could not be revealed by classical descriptors such as RMSD or sequence identity.

**Fig. 6.**
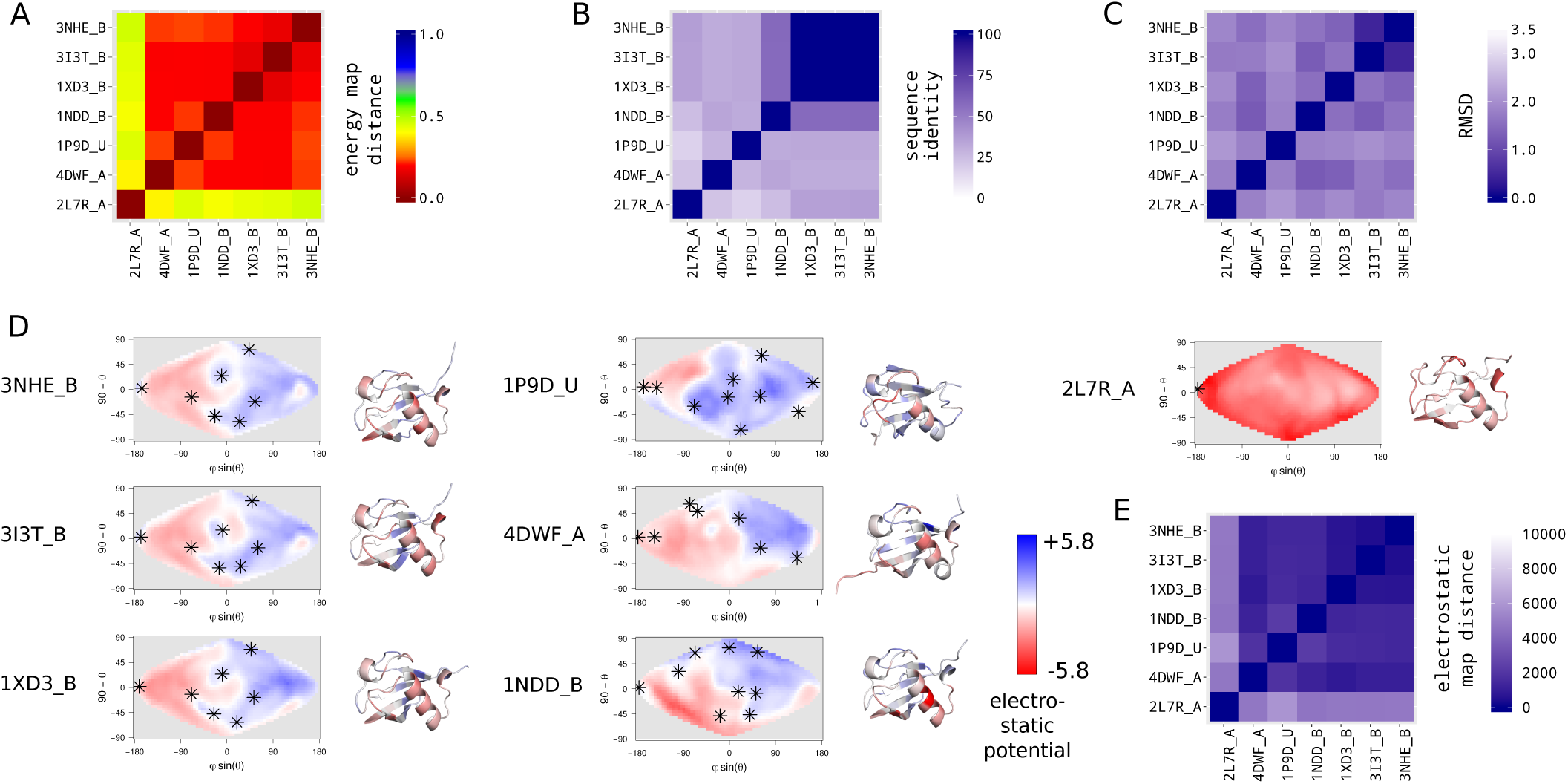
Ubiquitin-related family. (*A*) Energy map distances matrix. It corresponds to the subsection of the ADM for the ubiquitin-related family (for the construction of the ADM, see *Materials and Methods*). Each entry *(i,j)* represents the pairwise energy map distance of the ligand pair (*i,j*) averaged over the 74 receptors of the dataset. (*B*) Pairwise sequence identity matrix between all members of the family. (*C*) Pairwise root mean square deviation (RMSD) matrix between all members of the family. (*D*) Electrostatic maps and cartoon representations of the seven members of the family. An electrostatic map represents the distribution of the electrostatic potential on the surface of a protein (for more details, see S15 Fig and *Materials and Methods*). On the electrostatic maps, lysines positions are indicated by stars. Cartoon structures are colored according to the distribution of their electrostatic potential. (*E*) Electrostatic map distances matrix. Each entry *(i,j)* of the matrix represents the Manhattan distance between the electrostatic maps of the proteins *(i,j)*.

The AUC of 0.79 calculated previously with energy maps produced with the docking of either native-related or arbitrary pairs indicates that energy maps are specific to ligand families. To see whether this observation is not mainly due to the native-related pairs, we repeated the previous test while removing that time all energy maps computed with native-related pairs and calculated the resulting ADM. We then measured our ability to retrieve the homologs of each ligand by calculating the ROC curve as previously. The resulting AUC is still equal to 0.79, revealing that our ability to identify a ligand’s homologs is independent from the fact that the corresponding energy maps were computed with native-related or arbitrary pairs (Fig 5). This shows that the energy maps are specific to protein families whether the docked pairs are native-related or not. Consequently, the propensity of the whole protein surface to interact with a given ligand is conserved and specific to the ligand family whether the ligand is native-related or not. This striking result may reflect both positive and negative design operating on protein surfaces to maintain functional interactions and to limit random interactions that are inherent to a crowded environment.

### The interaction propensity of all surface regions of a receptor is evolutionary conserved for homologous ligands

To see whether some regions contribute more to the specificity of the maps produced by homologous ligands, we next dissected the effective contribution of the surface regions of the receptor defined according to their docking energy value, in the identification of ligand’s homologs. We discretized the energy values of each energy map into five categories, leading to a palette of five energy classes (see Fig 1D and *Materials and Methods*). These five-classes maps highlight low-energy regions (i.e. hot regions in red), intermediate-energy regions (i.e. warm, lukewarm and cool regions in orange, light-green and dark-green respectively) and high-energy regions (i.e. cold regions in blue). We first checked that the discretization of the energy maps does not affect our ability to identify the homologs of each of the 74 ligands from the comparison of their five-classes maps. The resulting AUC is 0.77 (Table 1), showing that the discretization step does not lead to an important loss of information.

**Table 1.** AUC obtained with different types of energy maps.

| type of<br>map | continuous<br>energy maps | five-classes<br>energy maps | hot energy<br>maps | warm energy<br>maps | lukewarm<br>energy maps | cool energy<br>maps | cold energy<br>maps |
| --- | --- | --- | --- | --- | --- | --- | --- |
| AUC | 0.79 | 0.77 | 0.73 | 0.76 | 0.76 | 0.76 | 0.79 |

Then, we evaluated the contribution of each of the five energy classes separately in the ligand’s homologs identification by testing our ability to retrieve the homologs of the 74 ligands from their one-class energy maps (either hot, warm, lukewarm, cool or cold) (see *Materials and Methods*). Table 1 shows the resulting AUCs. Interestingly, the information provided by each energy class taken separately is sufficient for discriminating the homologs of a given ligand from the rest of the dataset (Table 1). The resulting AUCs range from 0.76 to 0.79 for the warm, lukewarm, cool and cold classes and are comparable to those obtained with all classes taken together (0.77). This shows (i) that warm, lukewarm, cool, and cold regions alone are sufficient to retrieve homology relationships between ligands and (ii) that the localization on the receptor surface of a given energy class is specific to the ligand families. Hot regions are less discriminative and lead to an AUC of 0.73. In order to see how regions of an energy class are distributed over a receptor surface, we summed the one-class maps of the corresponding energy class calculated for this receptor into a stacked map (S16 Fig – see *Materials and Methods* for more details). A stacked map reflects the tendency of a surface region (i.e. map cells) to belong to the corresponding energy class. Fig 7 shows an example of the five stacked maps (i.e. for cold, cool, lukewarm, warm and hot regions) computed for the receptor 1P9D_U. Intermediates regions (i.e. warm, lukewarm and cool regions) are widespread on the stacked map while cold and hot regions are localized on few small spots (three and one respectively) no matter the nature of the ligand. S17 Fig shows for the receptor 1P9D_U the 12 cold and hot stacked maps computed for each ligand family separately. We can see that some cold spots are specific to ligand families and that their area distribution is specific to families while all 12 ligand families display the same hot spot in the map’s upper-right quadrant. These observations can be generalized to each receptor. On average, intermediate regions are widespread on the stacked maps and cover respectively 744, 1164 and 631 cells for cool, lukewarm and warm regions, while cold and hot regions cover no more than respectively 104 and 110 cells respectively (S18 Fig). Interestingly, hot regions are more colocalized than cold ones and are restricted to 2 distinct spots on average per stacked map, while cold regions are spread on 3.7 spots on average (t-Test *p =* 7.42e-13). These results show that ligands belonging to different families tend to dock preferentially on the same regions and thus lead to similar hot region distributions on the receptor surface. This observation recalls those made by *Fernandez-Recio et al*. [36], who showed that docking random proteins against a single receptor leads to an accumulation of low-energy solutions around the native interaction site and who suggested that different ligands will bind preferentially on the same localization.

**Fig. 7.**
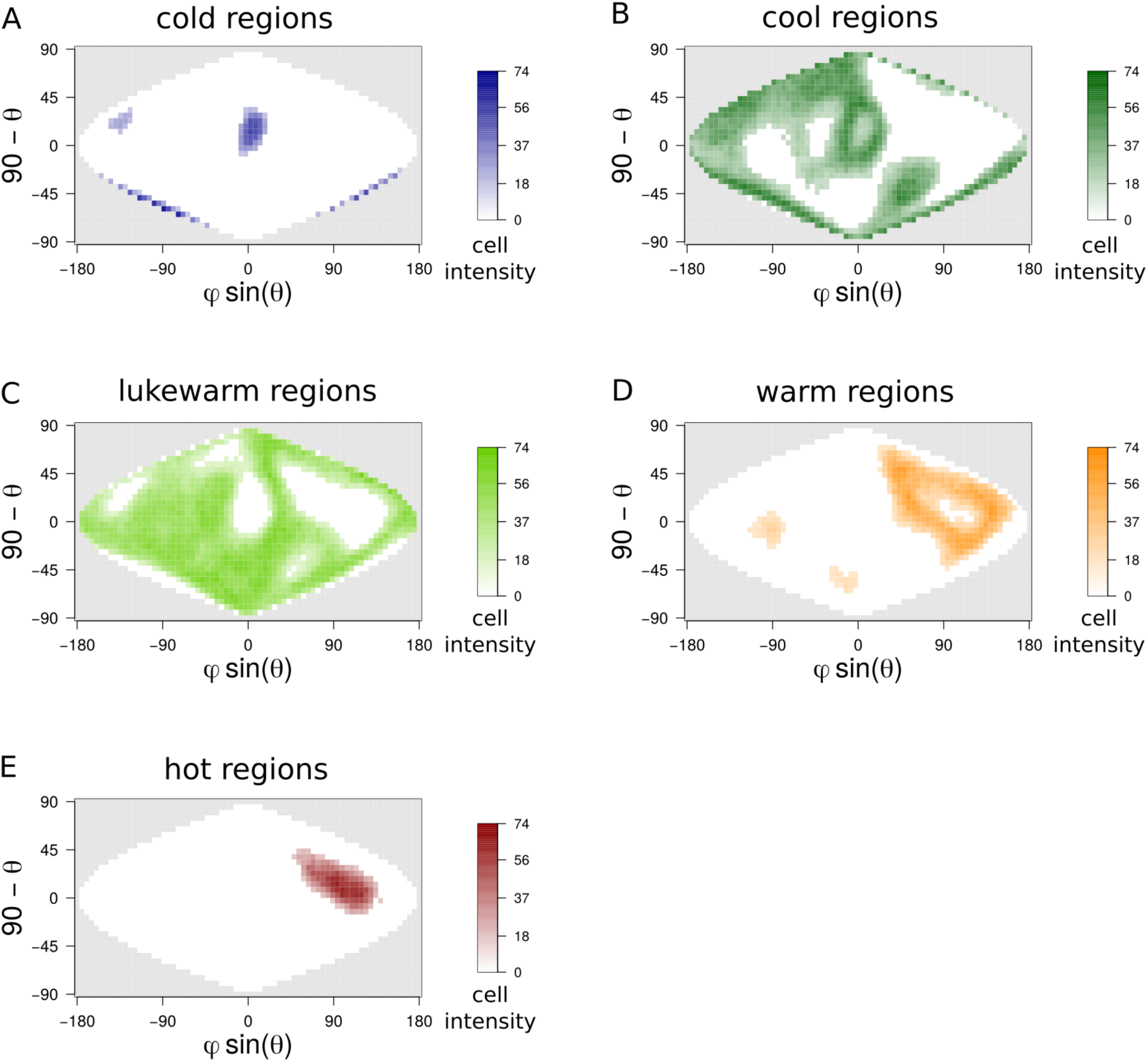
Stacked maps of 1P9D_U after the filtering of cells with too low intensity and areas of too small size. The protocol to generate stacked maps is presented in S16 Fig. (*A-E*) Stacked map for cold, cool, lukewarm, warm and hot regions respectively. The cell intensity in a stacked map of a given energy class indicates the number of times the cell has been associated to this energy class in all the corresponding one-class maps. One should notice that stacked maps of two different energy classes can overlap because a map cell can be associated to different energy classes depending on the docked ligands. S17 Fig presents cold and hot stacked maps of 1P9D_U computed for each ligand family.

We can hypothesize that hot regions present universal structural and biochemical features that make them more prone to interact with other proteins. To test this hypothesis, we computed for each protein of the dataset, the 2D projection of three protein surface descriptors (see *Materials and Methods* and S15 Fig): the Kyte-Doolittle (KD) hydrophobicity [44], the circular variance (CV) [45] and the stickiness [25]. The CV measures the density of protein around an atom and is a useful descriptor to reflect the local geometry of a surface region. CV values are comprised between 0 and 1. Low values reflect protruding residues and high values indicate residues located in cavities. Stickiness reflects the propensity of amino acids to be involved in protein-protein interfaces [25]. It is calculated as the log ratio of the residues frequencies on protein surfaces versus their frequencies in protein-protein interfaces. For each receptor, we calculated the correlation between the docking energy and the stickiness, hydrophobicity or CV over all cells of the corresponding 2D maps. We found a significant anti-correlation between the docking energy and these three descriptors (correlation test *p* between docking energies and respectively stickiness, hydrophobicity and CV < 2.2e-16, see S4 Table)). Fig 8 represents the boxplots of the stickiness, hydrophobicity and CV of each energy class (see S15 Fig and *Materials and Methods* section for more details). We observe a clear effect of these factors on the docking energy: cold regions are the less sticky, the less hydrophobic and the most protruding while hot ones are the most sticky, the most hydrophobic and the most planar (Tukey HSD test [46], *p* of the differences observed between each energy classes *<* 2.2e-16). One should notice that stickiness has been defined from a statistical analysis performed on experimentally characterized protein interfaces and therefore between presumed native partners. The fact that docking energies (physics-based) calculated either between native-related or arbitrary partners is anti-correlated with stickiness (statistics-based) defined from native interfaces, strengthens strongly the concept of stickiness as the propensity of interacting promiscuously and provides physics-based pieces of evidence for sticky regions as a proxy for promiscuous interactions.

**Fig. 8.**
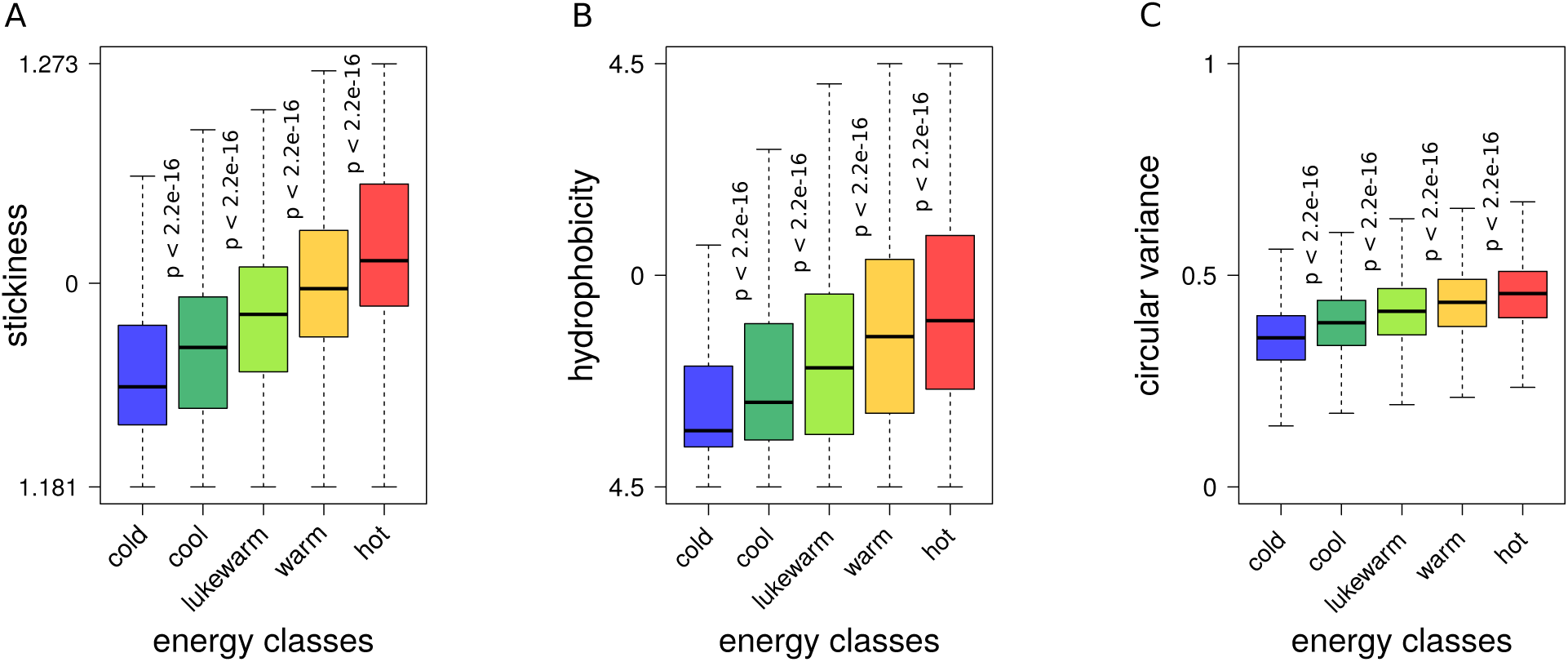
Boxplots of three descriptors of the protein surface. (*A*) the stickiness values, (*B*) the Kyte-Doolittle hydrophobicity and (*C*) the CV values, depending on the energy class. The stickiness, hydrophobicity and CV values are calculated for each protein following the protocol described in *Materials and Methods*. For each of these criteria, *p-values* between the median values of two “successive” energy classes were computed using the Tukey HSD statistical test [46].

We show that not only the area distribution on a receptor surface of hot regions but also those of intermediate and cold regions are similar for homologous ligands and are specific to ligand families (AUC ranging from 0.73 to 0.79) whether the ligands are native-related or not. This tendency is even stronger for intermediate and cold regions. Interestingly, the information contained in the cold regions that cover on average no more than 5.0% of the energy maps is sufficient to identify homology relationships between ligands.

## Discussion

In this study, we address the impact of both positive and negative design on thousands of interaction energy landscapes by the mean of a synthetic and efficient representation of the docking energy landscapes: two-dimensional energy maps that reflect the interaction propensity of the whole surface of a protein (namely the receptor) with a given partner (namely the ligand). We show that the distribution on the protein surface of all regions, including cold, intermediate and hot regions are similar for homologous ligands and are specific to ligand families whether the ligands are native-related or arbitrary. This reveals that the interaction propensity of the whole surface of proteins is constrained by functional and non-functional interactions, reflecting both positive and negative design operating on the whole surface of proteins, thus shaping the interaction energy landscapes of functional partners and random encounters. These observations were made on a dataset of 74 protein structures belonging to 12 families of structural homologs. 54 out of the 74 proteins of the dataset have at least one known partner in the dataset. For the 20 remaining proteins, we were not able to find evidences that they indeed interact with a protein of the dataset. However, we showed that the interaction propensity of a receptor is conserved for homologous ligands independently from the fact that these ligands correspond to native partners or not. Indeed, we showed that ligand homology relationships could be retrieved from their energy maps whether the maps were computed with native-related pairs or not (the corresponding AUCs calculated with and without native pairs both equal to 0.79).

Most studies that aim at depicting protein interactions focus on the functional ones and on the characterization of the native assembly modes [14,47–51]. Nevertheless, the importance of non-specific interactions and non-native assembly modes in protein interactions is no longer in doubt [7,19,21,27,52–55]. Experimental and *in-silico* studies showed the impact of non-specific interactions on the in-cell mobility of proteins [7,19,21,27]. In addition, an important literature describes the relationship between the physico-chemical properties of proteins and their ability for non-specific interactions [7,19,21,25,53]. In particular, Wang *et al* showed that the propensity for non-specific interactions is determined by multiple factors such as the protein charge, the conformational flexibility and the distribution of hydrophobic residues on the protein surface [19]. Finally, recent studies have demonstrated the importance of non-native assembly modes and non-interacting regions in the protein association process [54] and showed that it is relevant to consider them for predicting protein partners and binding affinities [56,57]. Particularly, Marin-Lopez *et al* developed a method based on the sampling of the conformational space of the encounter complexes formed during the binding process and showed that ΔG can be predicted accurately from the scoring of all encounter complexes sampled during a docking simulation, suggesting that the knowledge of the native pose is not necessary [57]. All these works highlight the importance of taking into account the whole surface of proteins as well as all the binding modes of a protein pair. This calls for the development of new methods that enable the systematic and physical characterization of the whole surface of a protein in interaction with a given partner. Here, we address the energy behavior of not only known binding sites, but also of the rest of the protein surface, which plays an important role in protein interactions by constantly competing with the native binding site. We show that the interaction propensity of the rest of the surface is not homogeneous and displays regions with different binding energies that are specific to ligand families. This may reflect the negative design operating on these regions to limit non-functional interactions [14,16,58]. We can hypothesize that non-interacting regions participate to favor functional assemblies (i.e. functional assembly modes with functional partners) over non-functional ones and are thus evolutionary constrained by non-functional assemblies. The fact that cold regions seem to be more specific to ligand families than hot ones may be explained by the fact that they are on average more protuberant and more charged. They thus display more variability than hot ones. Indeed, there is more variability in being positively or negatively charged and protuberant (with an important range of protuberant shapes) than in being neutral and flat. S19 Fig presents the electrostatic potential distribution of all energy classes. Cold regions display a larger variability of electrostatic potential (F-test, *p <* 2.2e-16) than hot regions that are mainly hydrophobic thus displaying neutral charge distributions in average. Consequently, a same hot region may be attractive for a large set of ligands while a cold region may be unfavorable to specific set of ligands, depending on their charges, shapes and other biophysical properties.

Moreover, we show that hot regions are very localized (4.9% of the cells of an energy map) and tend to be similar no matter the ligand. Similarly to protein interfaces that have been extensively characterized in previous studies [47,48,48–50], hot regions are likely to display universal properties of binding, i.e. they are more hydrophobic and more planar, and thus more “sticky” than the other regions. They may provide a non-specific binding patch that is suitable for many ligands. However, we can hypothesize that native partners have evolved to optimize their interfaces (positive design) so that native interactions prevail over non-native competing ones. Then positive design results in conserved binding sites and coevolved interfaces in order to maintain the charge and shape complementarity between functional partners. Indeed, we have previously shown that the docking of native partners lead to more favorable binding energies than the docking of non-native partners when the ligand is constrained to dock around the receptor’s native binding site [33,59]. All these results suggest a new physical model of protein surfaces where protein surface regions, in the crowded cellular environment, serve as a proxy for regulating the competition between functional and non-functional interactions. In this model, intermediate and cold regions play an important role by preventing non-functional assemblies and by guiding the interaction process towards functional ones and hot regions may select the functional assembly among the competing ones through optimized interfaces with the native partner. This model recalls the transitive model proposed by Marin-Lopez *et al* where a path connecting what they call “productive” (near-native) and “non-productive” (non-native) assemblies can be defined [57]. This path consists in distinct conformational states where each one is a macro-state of the binding process involving either the native binding site of each partner, a single native binding site or no native ones. The initial steps consist in macro-states which do not involve native binding sites. Macro-states appearing later during the assembly process would play a mechanistic role by drawing near the binding sites of the two partners. The latest stage would correspond to near-native conformations where van der Waals and de-solvation energies play a major role in the energy of interaction of the corresponding complexes while the electrostatic forces contribute mostly in the energy of non-native assemblies [60,61]. Figure S21 shows the effective electrostatic and van der Waals contributions in the total docking energy for the different surface regions (i.e. cold, intermediate and hot regions). Interestingly, our results concur with the observations made in [60,61] since we show that the contribution of electrostatic in the total docking energy is more important in cold regions while van der Waals energies predominate in hot ones. Characterizing the relationship between the macro-states defined by Marin-Lopez *et al* and the surface regions of different energy levels could provide at the same time a structural, physical and readable characterization of the binding process of two interacting proteins. In particular, it would be interesting to compare the properties of the different macro-states (involving or not the native binding sites of the two proteins) identified for functional and arbitrary pairs to see whether functional pairs displays specific features that would have resulted from an optimization of the binding process.

In this work, we used and extended the application of the 2D energy map representation developed in [36] to develop an original theoretical framework that enables the efficient, automated and integrative analysis of different protein surface features. Many other surface representations have been developed to characterize protein surface properties [62–67]. These representations include 2D projections or more sophisticated methods such as for example using 3D Zernike descriptors as a representation of the protein surface shape [68,69] which is a powerful tool to compare surface properties of either homologous or unrelated proteins since it does not require any prior alignment. 2D maps provide the area distribution of a given feature on the whole protein surface and their discretization enables the study of a given surface property (e.g. protuberance, planarity, stickiness, positively charged regions, or cold and hot regions for example). The advantage with the 2D energy maps is that they are easy to build and manipulate and their straightforward comparison enables (i) the study of relationships between different surface properties through the comparison of their area distributions on a protein surface and (ii) the highlight of the evolutionary constraints exerted on a given feature by comparing its area distribution on the surfaces of homologous proteins. Particularly, this enables the identification and characterization of hot regions on a protein surface which can be either specific or conserved for all ligands and opens up new possibilities for the development of novel methods for protein binding sites prediction and their classification as functional or promiscuous in the continuity of previous developments based on arbitrary docking [33,34,36,59].

Finally, our framework provides a proxy for further protein functional characterization as shown with the five proteins discussed in the *Results* section *Energy maps are specific to protein families*. The comparison of their respective energy maps enables us to reveal biophysical and functional properties that could not be revealed with classical monomeric descriptors such as RMSD or sequence identity. Indeed, our framework can reflect the energy behavior of a protein interacting with a subset of selected partners either functional or arbitrary, thus revealing functional and systemic properties of proteins. This work goes beyond the classical use of binary docking to provide a systemic point of view of protein interactions, for example by exploring the propensity of a protein to interact with hundreds of selected ligands, and thus addressing the behavior of a protein in a specific cellular environment. Particularly, exploring the dark interactome (i.e. non-functional assemblies and interactions with non-functional partners) can provide a wealth of valuable information to understand mechanisms driving and regulating protein-protein interactions. Precisely, our 2D energy maps based strategy enables its exploration in an efficient and automated way.

## Materials and Methods

### Protein dataset

The dataset comprises 74 protein structures divided into 12 families of structural homologs which were selected from the protein docking benchmark 5.0. (see S1 Table for a detailed list of each family). We decided to systematically remove all Antibody/Antigens complexes since they display specific evolutionary properties. Indeed, they did not co-evolve to interact and we can hypothesize that the evolutionary constraints operating on their interaction energy landscapes are different from those of other complexes. Each family is related to at least one other family (its native-related partners family) through a pair of interacting proteins for which the 3D structure of the complex is characterized experimentally (except the V set domain family: the two native partners are homologous and belong to the same family) (S1 Fig). Each family is composed of a monomer selected from the protein-protein docking benchmark 5.0 [70] in its bound and unbound forms, which is called the master protein. Each master protein has a native partner (for which the 3D structure of the corresponding complex has been characterized experimentally) in the database, which is the master protein for another family, except the V set domain family, which is a self-interacting family. When available, we completed families with interologs (i.e. pairs of proteins which have interacting homologs in an other organism) selected in the INTEREVOL database [71] according to the following criteria: (i) experimental structure resolution better than 3.25 Å, (ii) minimum alignment coverage of 75% with the rest of the family members and (iii) minimum sequence identity of 30% with at least one member of the family. Since we were limited by the number of available interologs, we completed families with unbound monomers homologous to the master following the same criteria and by searching for their partners in the following protein-protein interactions databases [72–77]. We consider that all members of a family correspond to native-related partners of all members of their native-related partner family. To address the impact of conformational changes of a protein on its interaction energy maps, we added different NMR conformers. We show that energy maps involving pairs of conformers are significantly more similar than those obtained for other pairs of homologous ligands (unilateral Wilcoxon test, *p* < 2.2e-16) showing that the conformational changes in a protein (lower than 3Å) have a low impact on the resulting energy maps (S20 Fig).

### Docking experiment and construction of energy maps

A complete cross-docking experiment was realized with the ATTRACT software [30] on the 74 proteins of the dataset, leading to 5476 (74 x 74) docking calculations (Fig 1A). ATTRACT uses a coarse-grain reduced protein representation and a simplified energy function comprising a pseudo Lennard-Jones term and an electrostatic term. The calculations took approximately 20000 hours on a 2.7GHz processor. Prior to docking calculations, all PDB structures were prepared with the DOCKPREP software [78].

During a docking calculation, the ligand L_i_ explores exhaustively the surface of the receptor R_k_ (whose position is fixed during the procedure), sampling and scoring thousands of different ligand docking poses (between 10000 and 50000 depending on the sizes of the proteins) (Fig 1A). For each protein couple R_k_-L_i_, a 2D energy map is computed which shows the distribution of the energies of all docking solutions over the receptor surface. To compute these maps, for all docking poses, the spherical coordinates (ϕ, θ) (with respect to the receptor center of mass (CM)) of the ligand CM are represented onto a 2D map in an equal-area 2D sinusoidal projection (Fig 1B) (see [36] for more details). Each couple of coordinates (ϕ, θ) is associated with the energy of the corresponding docking conformation (Fig 1B). A continuous energy map is then derived from the discrete one, where the map is divided into a grid of 36 x 72 cells. Each cell represents the same surface and, depending on the size of the receptor, can span from 2.5 Å^2^ to 13Å^2^. For each cell, all solutions with an energy score below 2.7 kcal/mol^−1^ from the lowest solution of the cell are retained, according to the conformations filtering protocol implemented in [33]. The average of the retained energy scores is then assigned to the cell. If there is no docking solution in a cell, a score of 0 is assigned to it. Finally, the energies of the cells are smoothed, by averaging the energy values of each cell and of the eight surrounding neighbors (Fig 1C).

For each map, the energy values are discretized into five energy classes of same range leading to a discrete five-colors energy map (Fig 1D). The range is calculated for each energy map and spans from the minimum to the maximum scores of the map cells. The range of the energy classes of the map R_k_-L_i_ is equal to (maxE – minE)/5, where maxE and minE correspond to the maximal and minimal energy values in the R_k_-L_i_ map. Each five-classes energy map is then split into five one-class maps, each one representing an energy class of the map (Fig 1E). The continuous, five-classes and one-class energy maps are calculated for the 5476 energy maps.

### Comparison of energy maps and identification of ligand’s homologs

Since, we cannot compare energy maps computed for two unrelated receptors, the procedure is receptor-centered and only compares energy maps produced with different ligands docked with the same receptor. The referential (i.e. the receptor) is thus the same (in other words all grid cells are comparable) for all the energy maps that are compared. For each receptor R_k_, we computed a 74×74 energy map distance (EMD) matrix where each entry (*i,j*) corresponds to the pairwise distance between the energy maps R_k_-L_i_ and R_k_-L_j_ resulting from the docking of the ligands L_i_ and L_j_ on the receptor R_k_ (Fig 3). The pairwise distance d_Man_(R_k_-L_i_, R_k_-L_j_) between the energy maps is calculated with a Manhattan distance according to equation (1)

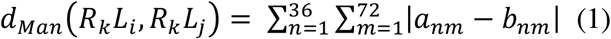

where a_nm_ and b_nm_ are the cells of row index *n* and column index *m* of the energy maps R_k_-L_i_ and R_k_-L_j_ respectively. Low distances reflect pairs of ligands that induce similar energy maps when they are docked on the same receptor. The procedure presented in Fig 3 is repeated for each receptor of the database resulting in 74 EMD matrices. The 74 EMD matrices are averaged into an averaged distances matrix (ADM). Each entry (*i,j*) of the ADM reflects the similarity of the R_k_-L_i_ and R_k_-L_j_ energy maps averaged over all the receptors R_k_ in the dataset. In order to estimate the extent to which family members display similar energy maps when they are docked with the same receptor, we tested our ability to correctly identify the homologs of the 74 ligands from the only comparison of its energy maps with those of the other ligands. Because, energy maps are receptor-centered, we cannot compare the energy maps computed for two unrelated receptors. The procedure consists in the comparison of energy maps produced with different ligands docked with a same receptor. Two ligands (*i,j*) are predicted as homologs according to their corresponding distance (*i,j*) in the ADM. Values close to zero should reflect homologous ligand pairs, while values close to one should reflect unrelated ligand pairs. A Receiver Operating Characteristic (ROC) curve and its Area Under the Curve (AUC) are computed from the ADM. True positives (TP) are all the homologous ligand pairs and predicted as such, true negatives (TN) are all the unrelated ligand pairs and predicted as such. False positives (FP) are unrelated ligand pairs but incorrectly predicted as homologous pairs. False negatives (FN) are homologous ligand pairs but incorrectly predicted as unrelated pairs. ROC curves and AUC values were calculated with the R package pROC [79]. The ligand’s homologs identification was also realized using the five-classes energy maps or the one-class energy maps taken separately. The five energy class regions display very different sizes, with median ranging from 63 and 66 cells for the cold and hot regions to 633 cells for the yellow one. To prevent any bias due to the size of the different classes, we normalized the Manhattan distance by the size of the regions compared in the map. The rest of the procedure is the same than those used for continuous energy maps (Fig 3).

To visualize the area distribution of the regions of a given energy class for all ligands on the receptor surface, the 74 corresponding one-class maps are summed into a stacked map where each cell’s intensity varies from 0 to 74 (S16 Fig). To remove background-image from these maps, i.e. cells with low intensity (intensity < 17) and the areas of small size (< 4 cells), we used a Dirichlet process mixture model simulation for image segmentation (R package *dpmixsim*) [80].

### 2D projection of monomeric descriptors of protein surfaces

We computed KD hydrophobicity [44], stickiness [25], CV [45] maps of each protein of the dataset, in order to compare their topology with the energy maps. Prior to all, proteins belonging to the same families were structurally aligned with TM-align [81] in order to place them in the same reference frame, making their maps comparable. Particles were generated around the protein surface with a slightly modified Shrake-Rupley algorithm [82]. The density of spheres is fixed at 1Å^2^, representing several thousands particles per protein. Each particle is located at 5Å from the surface of the protein. The CV, stickiness and KD hydrophobicity values of the closest atom of the protein are attributed to each particle. We also generated electrostatic maps reflecting the distribution of the contribution of the electrostatic potential on a protein surface. The electrostatics potential was computed with the APBS software suite [38] using the CHARMM force field [83]. In this case the procedure is different as the electrostatic potential is calculated at each particle position, using the multivalue executable from the APBS software suite.

The CV was calculated following the protocol described in [45] on the all-atom structures. Stickiness and hydrophobicity were calculated on ATTRACT coarse-grain models. After attributing a value to each particle, the position of their spherical coordinates is represented in a 2-D sinusoidal projection, following the same protocol as described in Fig 1 and *Materials and Methods* section *Docking experiment and construction of energy maps*. The map is then smoothed following the protocol in Fig 1.

## Acknowledgments

We thank F. Fraternali, R. Guerois, E. Laine, and M. Montes for their constructive comments on the manuscript.

## Notes

The authors have declared that no competing interests exist.

