## Supplementary Material file for "Protein interaction energy landscapes are shaped by functional and also non-functional partners"

**S1 Table. List of proteins of the dataset and their structural families.**

| Protein identifier | sequence identity (%) | RMSD (Å$^2$) | Structural family |
|---|---|---|---|
| 1YVB_A (b)<br>2GHU_A (u)<br>2NQD_B (b)<br>3F75_A (b)<br>3BWK_C (b)<br>3RVV_A (b)<br>1YAL_B (u)<br>3IMA_A (b) | 43 | 1.57 | **Papain-like** |
| 1QA9_A (b)<br>1CCZ_A (u)<br>1QA9_B (b)<br>1HNF_A (u)<br>2PTT_A (b)<br>2PTT_B (b) | 33.6 | 1.74 | **V set domains (antibody variable domain-like)** |
| 1XD3_A (b)<br>1UCH_A (u)<br>1CMX_A (b)<br>2WDT_A (b)<br>3IFW_A (b) | 45.8 | 1.71 | **Ubiquitin carboxyl-terminal hydrolases (UCH-L)** |
| 2AYO_A (b)<br>2AYN_A (u)<br>3I3T_A (b)<br>3NHE_A (b)<br>2Y6E_A (u) | 41.1 | 2.35 | **Ubiquitin carboxyl-terminal hydrolases (UCH)** |
| 3FN1_A (b)<br>2LQ7_A_2 (u)<br>2LQ7_A_13 (u)<br>2LQ7_A_15 (u)<br>2LQ7_A_19 (u) | 100 | 2.13 | **Ubiquitin activating enzymes (UBA)** |
| 3FN1_B (b)<br>2EDI_A_1 (u)<br>2EDI_A_7 (u)<br>2EDI_A_10 (u)<br>1YH2_A (u)<br>1FXT_A (b)<br>2GMI_A (b)<br>4P5O_G (b) | 48 | 1.79 | **UBC-related** |
| 1AVW_B (b)<br>1BA7_A (u)<br>3I2A_B (u)<br>3BX1_B (b)<br>4AN6_A (u)<br>4J2Y_A (b)<br>4IHZ_A (u) | 37.1 | 2.22 | **Kunitz (STI) inhibitors** |
| 1M9X_B (b)<br>4J93_A (u)<br>2WLV_B (u)<br>2XGU_A (u) | 52.7 | 1.93 | **Retrovirus capsid proteins, N-terminal core domain** |
| 1XD3_B (b)<br>3NHE_B (b)<br>3I3T_B (b)<br>1NDD_B (u)<br>2L7R_A (u)<br>1P9D_U (u)<br>4DWF_A (u) | 44.2 | 1.64 | **Ubiquitin-related** |
| 1YVB_B (b)<br>1CEW_I (u)<br>3IMA_B (b)<br>4N6O_B (b)<br>2CH9_B (u) | 40 | 1.99 | **Cystatins** |
| 1M9X_A (b)<br>2CPL_A (u)<br>3K2C_A (u)<br>1MZW_A (u)<br>2RMC_G (u)<br>2NUL_B (u) | 52.2 | 1.53 | **Cyclophilins (peptidylprolyl isomerase)** |
| 1AVW_A (b)<br>1QQU_A (u)<br>1FXY_B (u)<br>1HNE_C (u)<br>1ZJD_A (b)<br>1BZX_A (b)<br>4BNR_A (b)<br>3I29_A (b) | 49.6 | 1.45 | **Eukaryotic proteases** |

Proteins are referred by their PDB identifiers, followed by their chain identifier. The NMR conformers are referred with their conformation identifier. The conformational state of the structures are indicated in brackets ((b) for bound conformation, (u) for unbound conformation). Structural families are named according to the SCOPe database [1] at the family level. Averaged sequence identity and RMSD are given for each family.

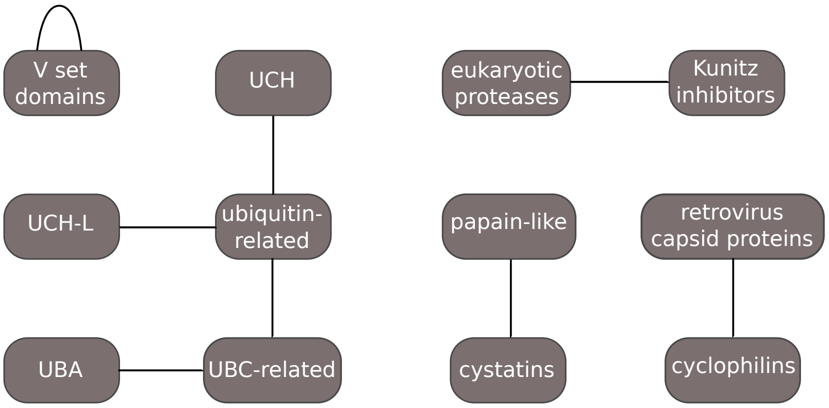


**S1 Fig. Interactions between structural families of the dataset.** Interactions are symbolized by links between families. An interaction is established between two families when, there is at least one PDB reporting a structure of complex involving members of the two families [2]. Consequently, all members of a family do not necessarily have its native partner in its native-related partner family. The V set domains family is a special case of self-interacting family, where members form dimers of structural homologs.

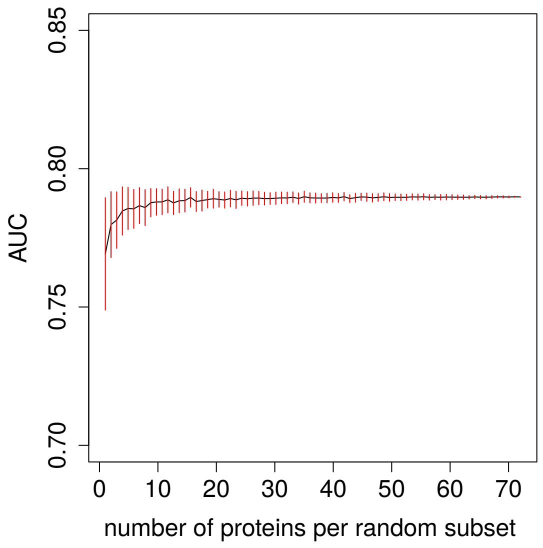


**S2 Fig. AUC values calculated on random subsets of receptor of different sizes.** The AUC is computed following the protocol described in Fig. 1 with random subsets composed from 1 to 73 receptors. Receptors of each subset are randomly chosen among the 74 receptors of the dataset. For each subset size, the procedure is repeated 100 times. Red vertical lines indicate the standard deviation of the AUC for each subset size. Above a subset size of five receptors, the AUC does not significantly fluctuate (risk of wrongly rejecting the equality of two variances (F-test) >5% [3]).

**S2 Table. AUC according to the grid resolution used for the energy maps**

| grid resolution | 144x72 | 120x60 | 100x50 | 80x40 | 72x36 | 60x30 | 48x24 |
|---|---|---|---|---|---|---|---|
| AUC | 0.8 | 0.8 | 0.8 | 0.79 | 0.79 | 0.78 | 0.78 |

The grid resolution corresponds to the number of cells composing the energy maps. The AUC is calculated following the same protocol used in the main text (see *Materials and Methods*)

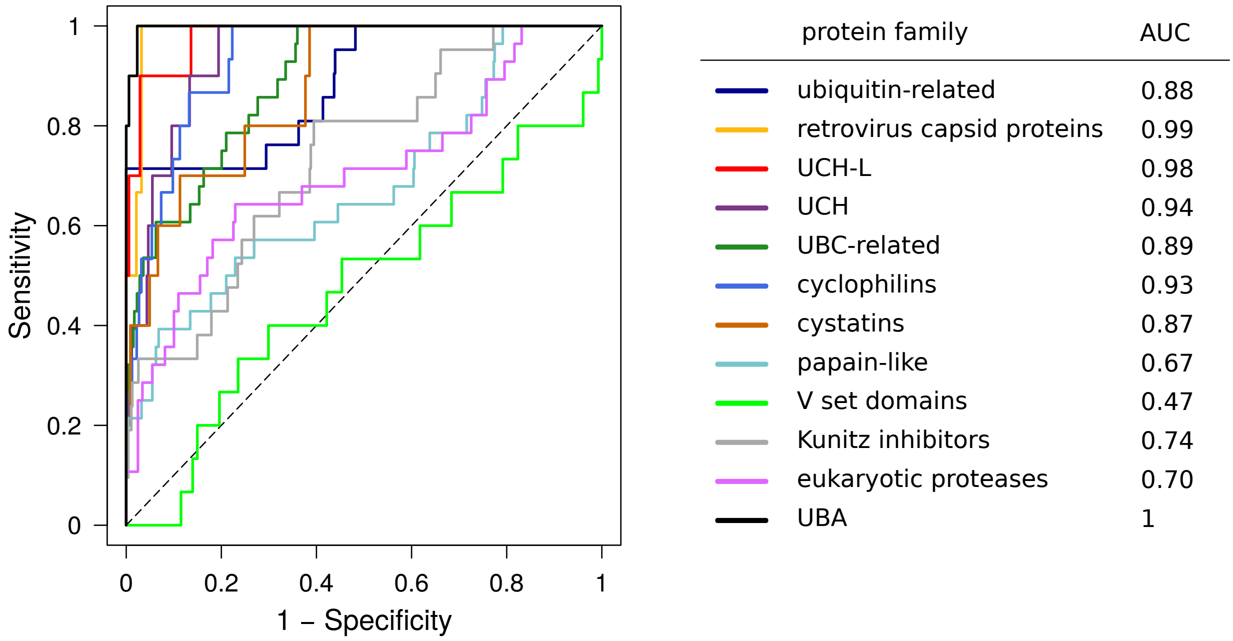


**S3 Fig. Receiver operating characteristic (ROC) curve and Area Under this Curve (AUC) calculated for each family.**

**S3 Table. Estimation of the effective contribution of sequence identity, RMSD and electrostatic distance in the pairwise ADM distances for each ligand pair belonging to a same family**

| protein family | sequence identity | RMSD | electrostatics potential |
|---|---|---|---|
| ubiquitin-related | 0.02396 | 0.01164 | 4.228e-5 |
| V set domains | 0.00651 | 0.5359 | 0.1773 |
| UCH-L | 0.07388 | 0.1076 | 0.3466 |
| UCH | 0.5109 | 0.00651 | 0.3104 |
| UBA | NA | 0.00117 | 0.04791 |
| UBC-related | 0.00566 | 0.3972 | 0.01296 |
| Kunitz inhibitors | 0.00527 | 0.00358 | 2.989e-5 |
| retrovirus capsid proteins | 0.00481 | 0.04156 | 0.00481 |
| papain-like | 0.8075 | 0.4613 | 9.09e-7 |
| cystatins | 0.0706 | 0.9867 | 0.08972 |
| cyclophilins | 0.03993 | 0.2539 | 0.00028 |
| eukaryotic proteases | 0.3861 | 0.2273 | 5.634e-7 |
| all proteins | 0.64989 | 0.91385 | 2.59e-7 |

A linear model was constructed from the dataset constituted of all the intra-family ligand pairs (202 protein pairs). This model allows the estimation of the linear correlation between the three descriptors and the pairwise ADM distance. The model takes into account the individual contribution of each descriptor as well as their crossed contributions with each other. The p-value of each individual contribution calculated over the 202 pairs is estimated with a Fisher test and are given in the table line “all proteins”. We then individually looked each family to see whether the contribution of the descriptors is dependent from the family. Inside each family, the number of protein pairs is too small to estimate a linear model. Consequently, we used a Spearman correlation coefficient test to estimate the p-value of each contribution.

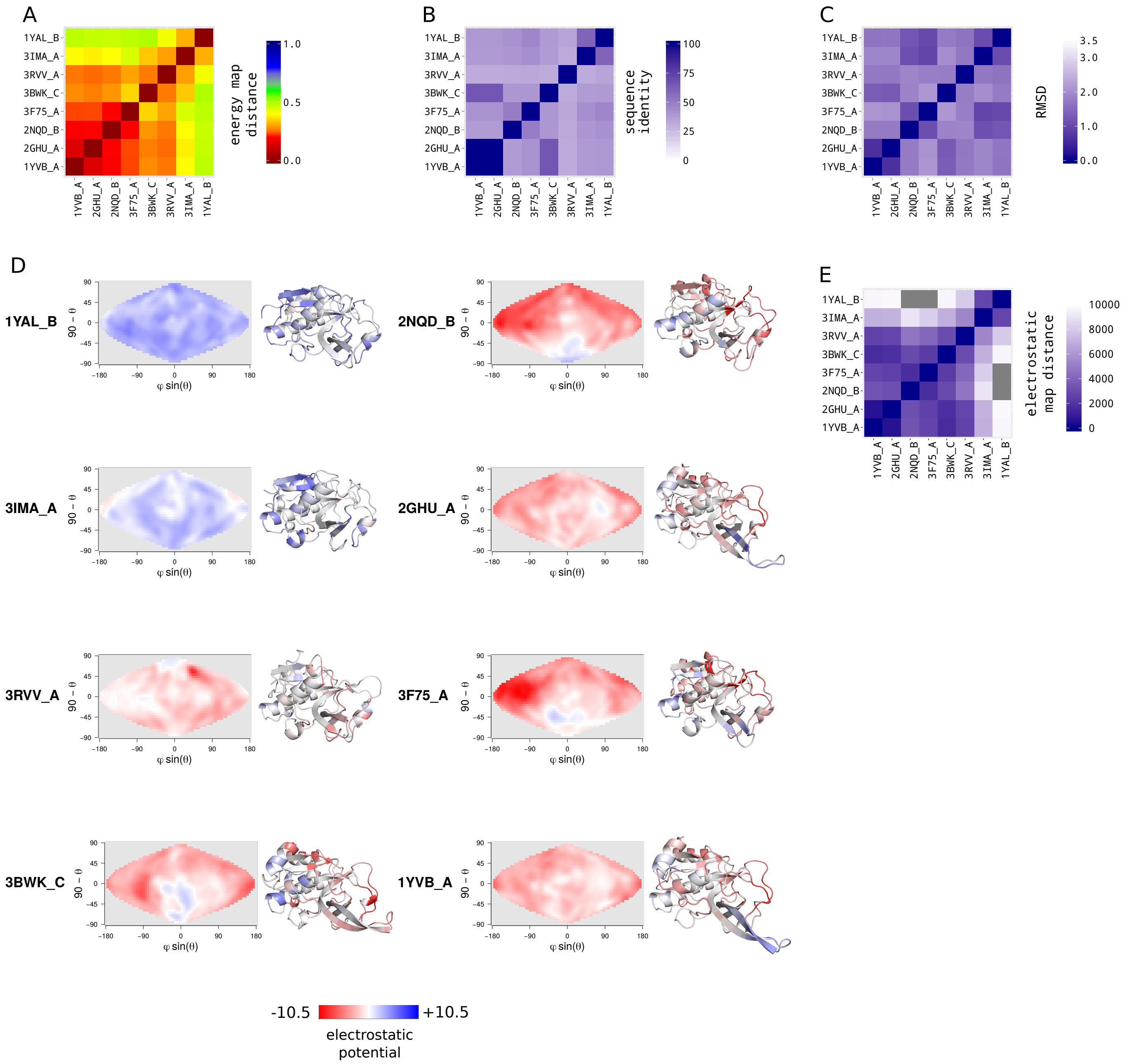


**S4 Fig. Papain-like family.** (*A*) Energy map distances matrix. It corresponds to the subsection of the ADM for the papain-like family (for the construction of the ADM, see *Materials and Methods*). Each entry *(i,j)* represents the pairwise energy map distance of the ligand pair *(i,j)* averaged over the 74 receptors of the dataset (for more details, see *Materials and Methods*). (*B*) Pairwise sequence identity matrix between all members of the family. (*C*) Pairwise root mean square deviation (RMSD) matrix between all members of the family. (*D*) Electrostatic maps and cartoon representations of the seven members of the family. An electrostatic map represents the distribution of the electrostatic potential on the surface of a protein (see Fig. S15 and *Materials and Methods*). Cartoon structures are colored according to the distribution of their electrostatic potential. (*E*) Electrostatic map distances matrix. Each entry *(i,j)* of the matrix represents the Manhattan distance between the electrostatic maps of the proteins *(i,j)*.

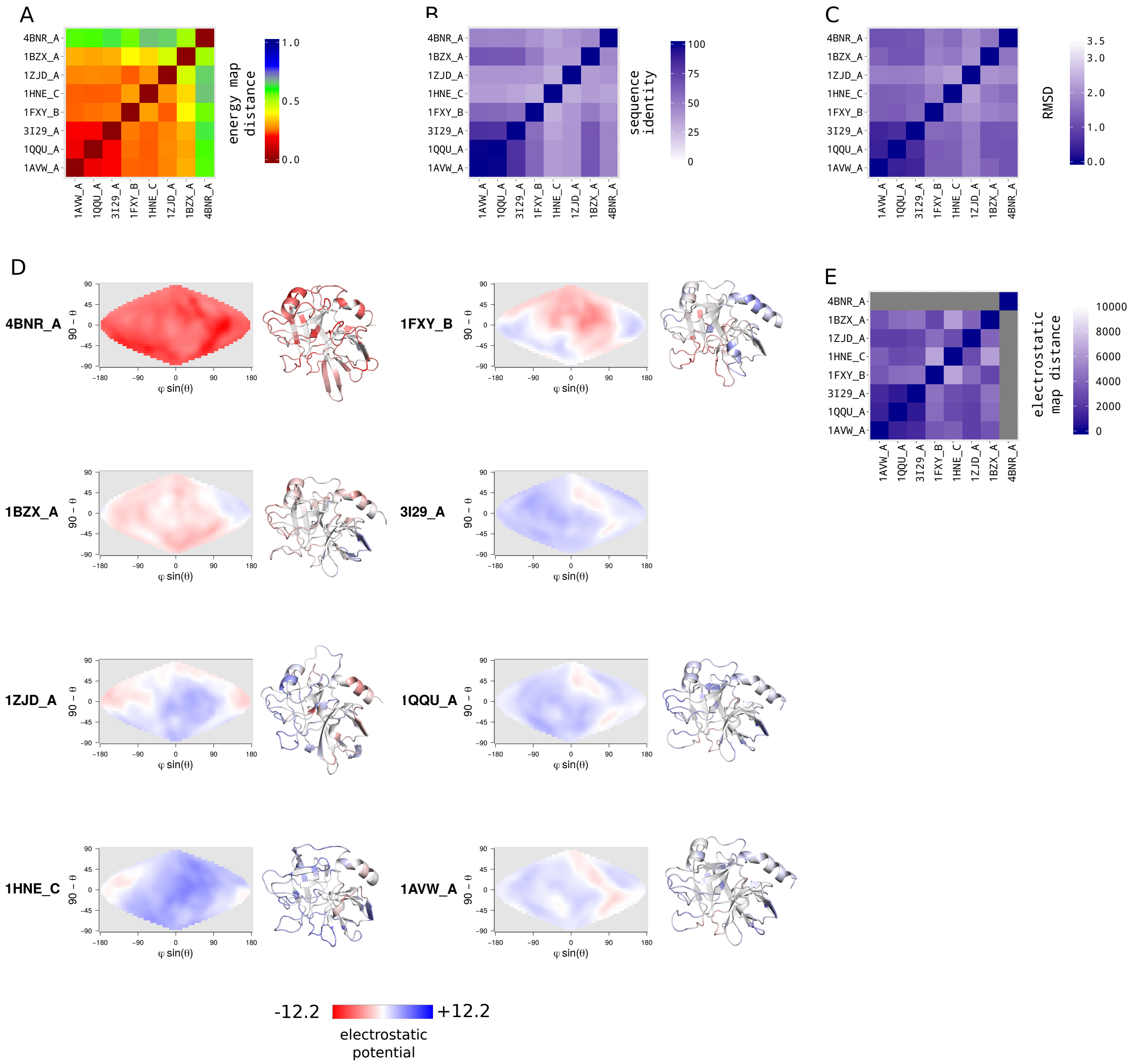


**S5 Fig. Eukaryotic-proteases family.** (*A*) Energy map distances matrix. It corresponds to the subsection of the ADM for the Eukaryotic proteases family (for the construction of the ADM, see *Materials and Methods*). Each entry *(i,j)* represents the pairwise energy map distance of the ligand pair *(i,j)* averaged over the 74 receptors of the dataset (for more details, see *Materials and Methods*). (*B*) Pairwise sequence identity matrix between all members of the family. (*C*) Pairwise root mean square deviation (RMSD) matrix between all members of the family. (*D*) Electrostatic maps and cartoon representations of the seven members of the family. An electrostatic map represents the distribution of the electrostatic potential on the surface of a protein (for more details, see Fig. S15 and *Materials and Methods*). Cartoon structures are colored according to the distribution of their electrostatic potential. (*E*) Electrostatic map distances matrix. Each entry *(i,j)* of the matrix represents the Manhattan distance between the electrostatic maps of the proteins *(i,j)*.

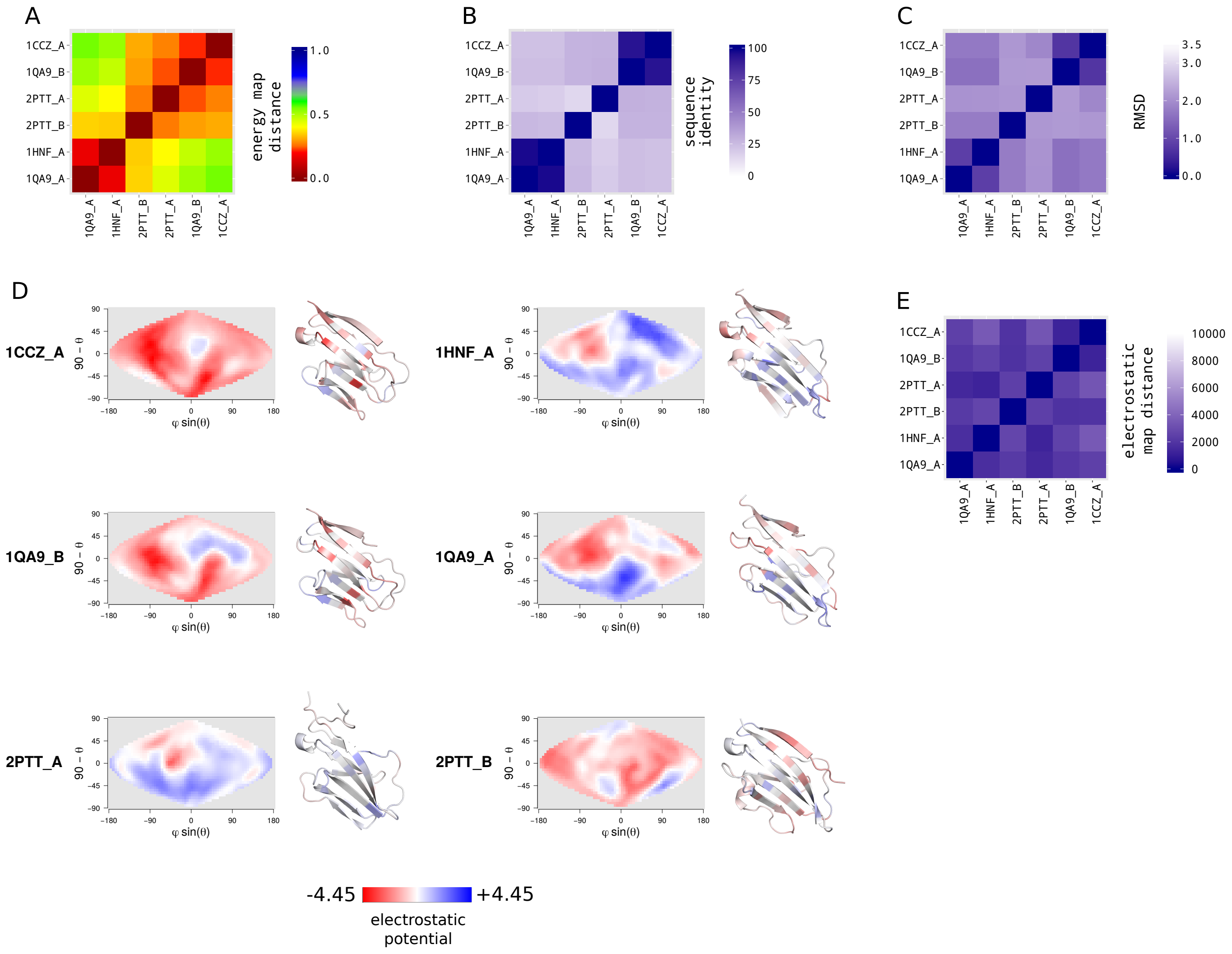


**S6 Fig. V set domains family.** (*A*) Energy map distances matrix. It corresponds to the subsection of the ADM for the V set domain family (for the construction of the ADM, see *Materials and Methods*). Each entry *(i,j)* represents the pairwise energy map distance of the ligand pair *(i,j)* averaged over the 74 receptors of the dataset (for more details, see *Materials and Methods*). (*B*) Pairwise sequence identity matrix between all members of the family. (*C*) Pairwise root mean square deviation (RMSD) matrix between all members of the family. (*D*) Electrostatic maps and cartoon representations of the six members of the family. An electrostatic map represents the distribution of the electrostatic potential on the surface of a protein (for more details, see Fig. S15 and *Materials and Methods*). Cartoon structures are colored according to the distribution of their electrostatic potential. (*E*) Electrostatic map distances matrix. Each entry *(i,j)* of the matrix represents the Manhattan distance between the electrostatic maps of the proteins *(i,j)*.

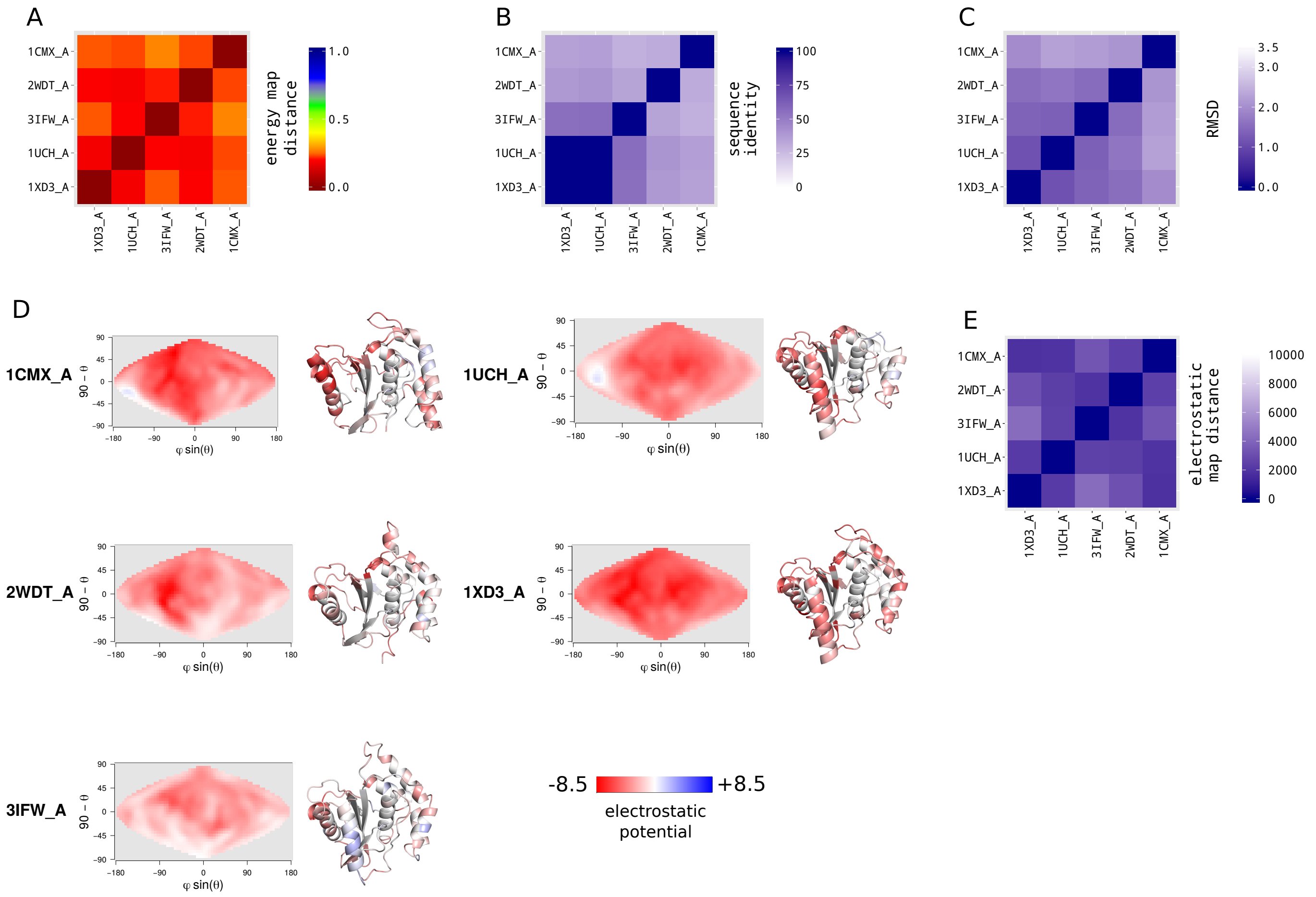


**S7 Fig. UCH-L family.** (*A*) Energy map distances matrix. It corresponds to the subsection of the ADM for the UCH-L family (for the construction of the ADM, see *Materials and Methods*). Each entry *(i,j)* represents the pairwise energy map distance of the ligand pair *(i,j)* averaged over the 74 receptors of the dataset (for more details, see *Materials and Methods*). (*B*) Pairwise sequence identity matrix between all members of the family. (*C*) Pairwise root mean square deviation (RMSD) matrix between all members of the family. (*D*) Electrostatic maps and cartoon representations of the seven members of the family. An electrostatic map represents the distribution of the electrostatic potential on the surface of a protein (for more details, see Fig. S15 and *Materials and Methods*). Cartoon structures are colored according to the distribution of their electrostatic potential. (*E*) Electrostatic map distances matrix. Each entry *(i,j)* of the matrix represents the Manhattan distance between the electrostatic maps of the proteins *(i,j)*.

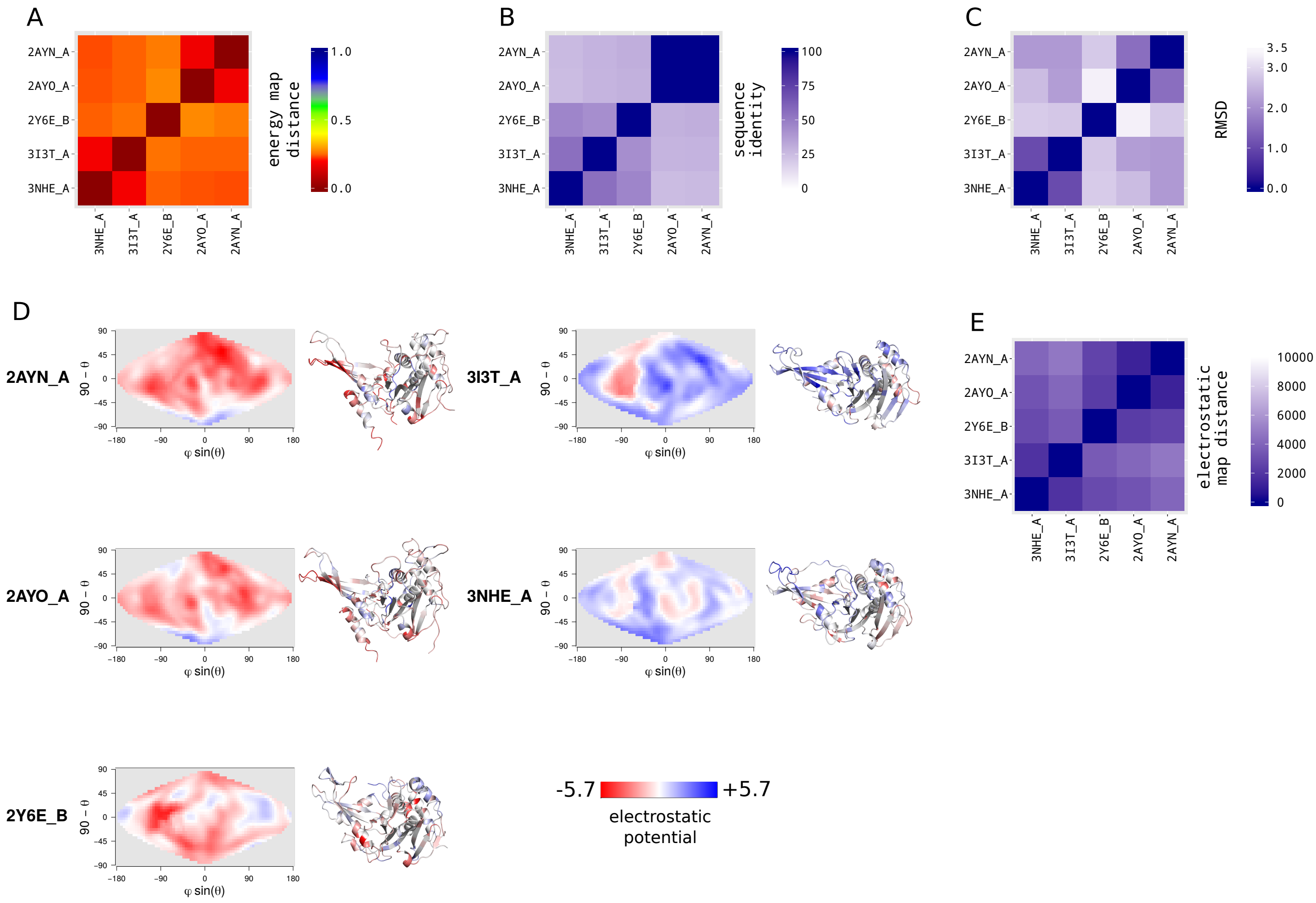


**S8 Fig. UCH family.** (*A*) Energy map distances matrix. It corresponds to the subsection of the ADM for the UCH family (for the construction of the ADM, see *Materials and Methods*). Each entry *(i,j)* represents the pairwise energy map distance of the ligand pair *(i,j)* averaged over the 74 receptors of the dataset (for more details, see *Materials and Methods*). (*B*) Pairwise sequence identity matrix between all members of the family. (*C*) Pairwise root mean square deviation (RMSD) matrix between all members of the family. (*D*) Electrostatic maps and cartoon representations of the seven members of the family. An electrostatic map represents the distribution of the electrostatic potential on the surface of a protein (for more details, see Fig. S15 and *Materials and Methods*). Cartoon structures are colored according to the distribution of their electrostatic potential. (*E*) Electrostatic map distances matrix. Each entry *(i,j)* of the matrix represents the Manhattan distance between the electrostatic maps of the proteins *(i,j)*.

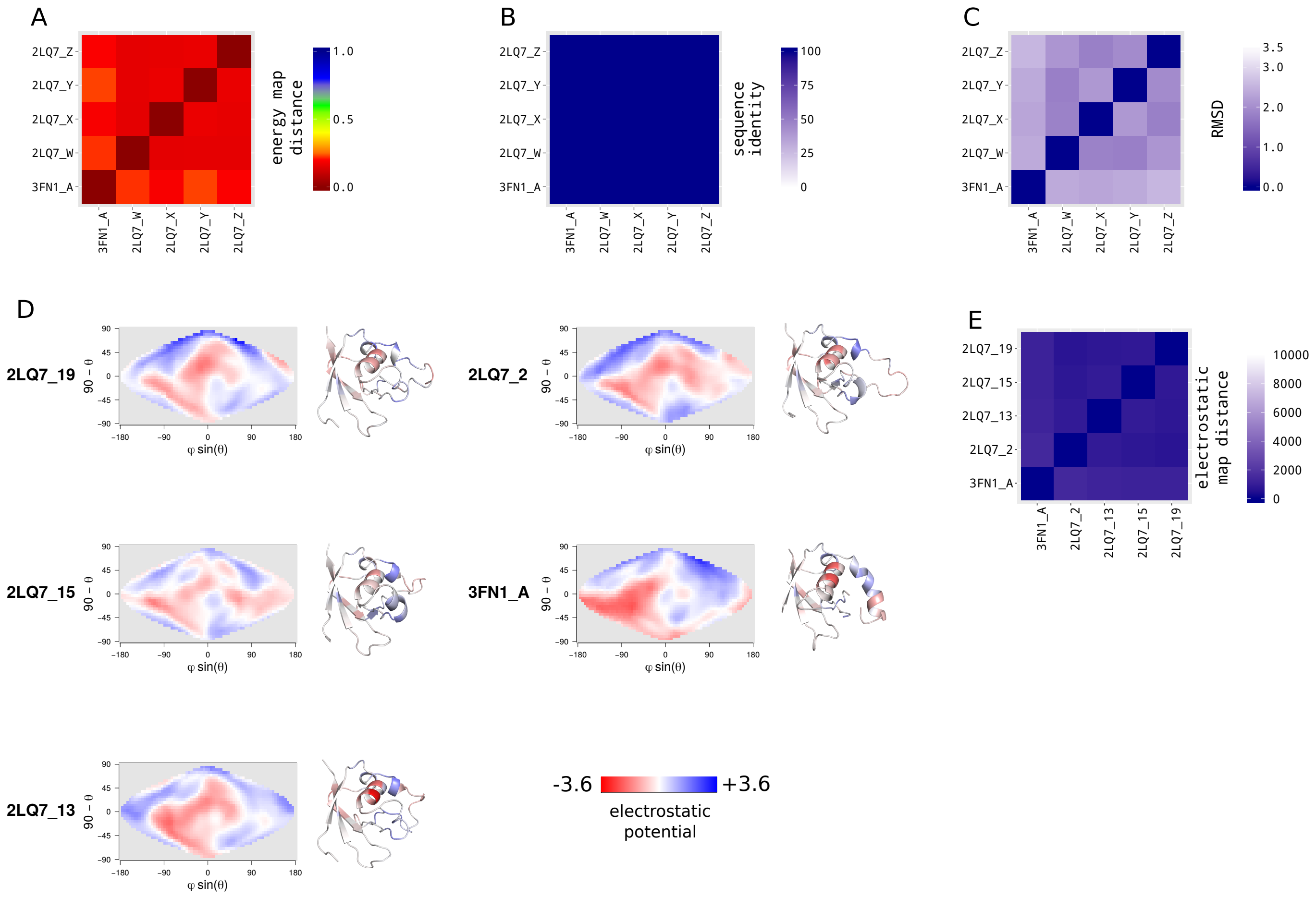


**S9 Fig. Ubiquitin activating enzymes family.** (*A*) Energy map distances matrix. It corresponds to the subsection of the ADM for the Ubiquitin activating enzymes family (for the construction of the ADM, see *Materials and Methods*). Each entry *(i,j)* represents the pairwise energy map distance of the ligand pair *(i,j)* averaged over the 74 receptors of the dataset (for more details, see *Materials and Methods*). (*B*) Pairwise sequence identity matrix between all members of the family. (*C*) Pairwise root mean square deviation (RMSD) matrix between all members of the family. (*D*) Electrostatic maps and cartoon representations of the seven members of the family. An electrostatic map represents the distribution of the electrostatic potential on the surface of a protein (for more details, see Fig. S15 and *Materials and Methods*). Cartoon structures are colored according to the distribution of their electrostatic potential. (*E*) Electrostatic map distances matrix. Each entry *(i,j)* of the matrix represents the Manhattan distance between the electrostatic maps of the proteins *(i,j)*.

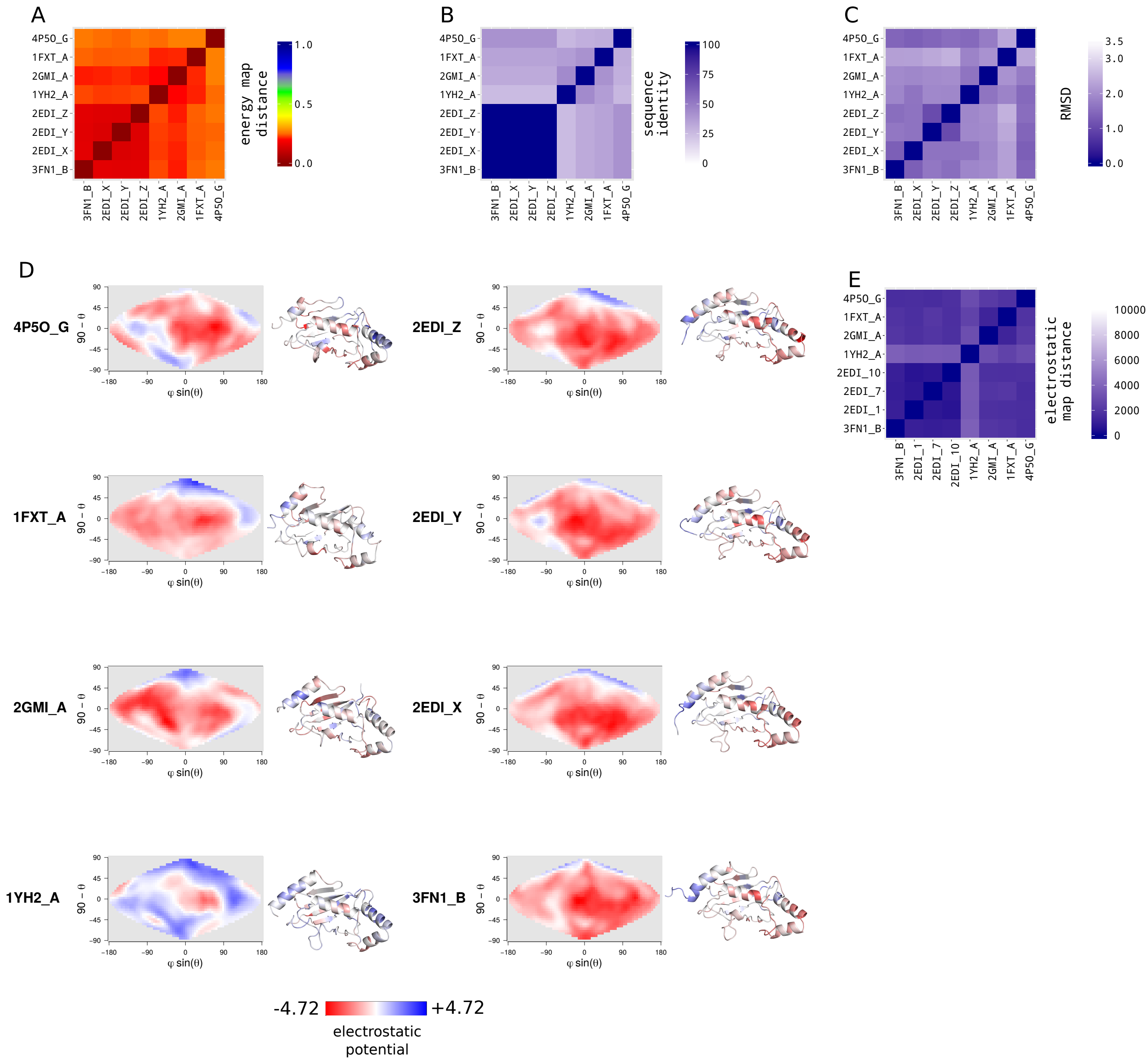


**S10 Fig. UBC-related family.** (*A*) Energy map distances matrix. It corresponds to the subsection of the ADM for the UBC-related family (for the construction of the ADM, see *Materials and Methods*). Each entry *(i,j)* represents the pairwise energy map distance of the ligand pair *(i,j)* averaged over the 74 receptors of the dataset (for more details, see *Materials and Methods*). (*B*) Pairwise sequence identity matrix between all members of the family. (*C*) Pairwise root mean square deviation (RMSD) matrix between all members of the family. (*D*) Electrostatic maps and cartoon representations of the seven members of the family. An electrostatic map represents the distribution of the electrostatic potential on the surface of a protein (for more details, see Fig. S15 and *Materials and Methods*). Cartoon structures are colored according to the distribution of their electrostatic potential. (*E*) Electrostatic map distances matrix. Each entry *(i,j)* of the matrix represents the Manhattan distance between the electrostatic maps of the proteins *(i,j)*.

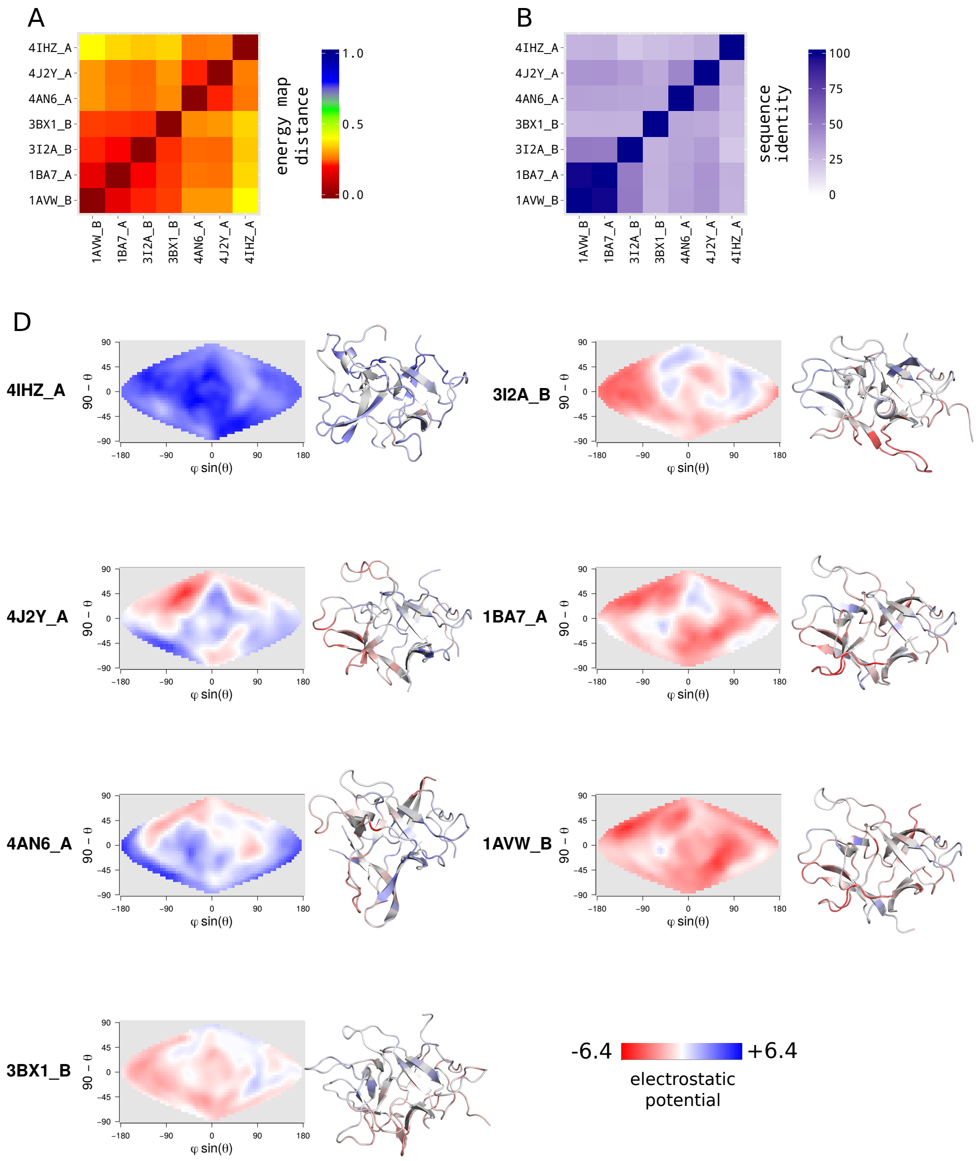


**S11 Fig. Kunitz (STI) inhibitors family.** (*A*) Energy map distances matrix. It corresponds to the subsection of the ADM for the Kunitz (STI) inhibitors family (for the construction of the ADM, see *Materials and Methods*). Each entry *(i,j)* represents the pairwise energy map distance of the ligand pair *(i,j)* averaged over the 74 receptors of the dataset (for more details, see *Materials and Methods*). (*B*) Pairwise sequence identity matrix between all members of the family. (*C*) Pairwise root mean square deviation (RMSD) matrix between all members of the family. (*D*) Electrostatic maps and cartoon representations of the seven members of the family. An electrostatic map represents the distribution of the electrostatic potential on the surface of a protein (for more details, see Fig. S15 and *Materials and Methods*). Cartoon structures are colored according to the distribution of their electrostatic potential. (*E*) Electrostatic map distances matrix. Each entry *(i,j)* of the matrix represents the Manhattan distance between the electrostatic maps of the proteins *(i,j)*.

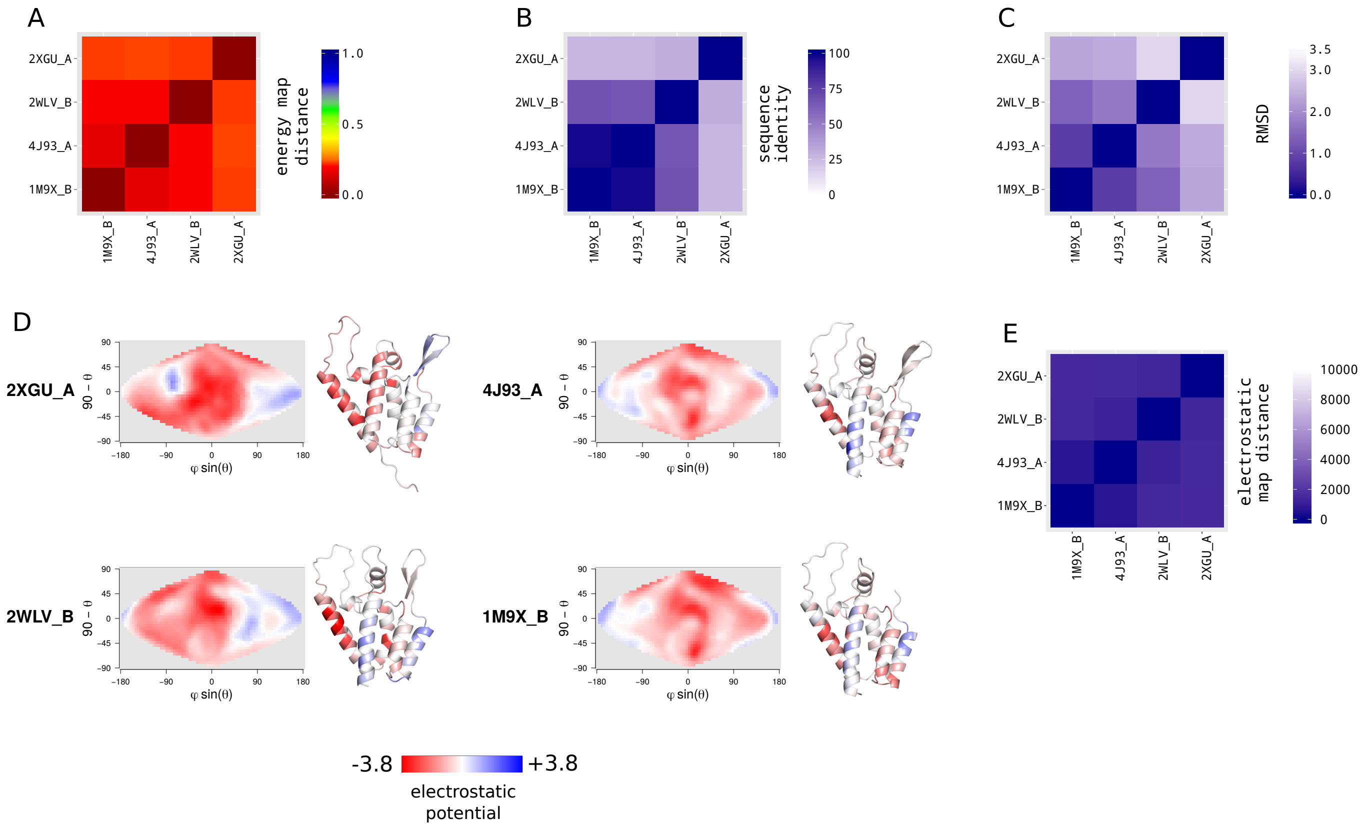


**S12 Fig. Retrovirus capsid proteins family.** (*A*) Energy map distances matrix. It corresponds to the subsection of the ADM for the retrovirus capsid proteins family (for the construction of the ADM, see *Materials and Methods*). Each entry *(i,j)* represents the pairwise energy map distance of the ligand pair *(i,j)* averaged over the 74 receptors of the dataset (for more details, see *Materials and Methods*). (*B*) Pairwise sequence identity matrix between all members of the family. (*C*) Pairwise root mean square deviation (RMSD) matrix between all members of the family. (*D*) Electrostatic maps and cartoon representations of the seven members of the family. An electrostatic map represents the distribution of the electrostatic potential on the surface of a protein (for more details, see Fig. S15 and *Materials and Methods*). Cartoon structures are colored according to the distribution of their electrostatic potential. (*E*) Electrostatic map distances matrix. Each entry *(i,j)* of the matrix represents the Manhattan distance between the electrostatic maps of the proteins *(i,j)*.

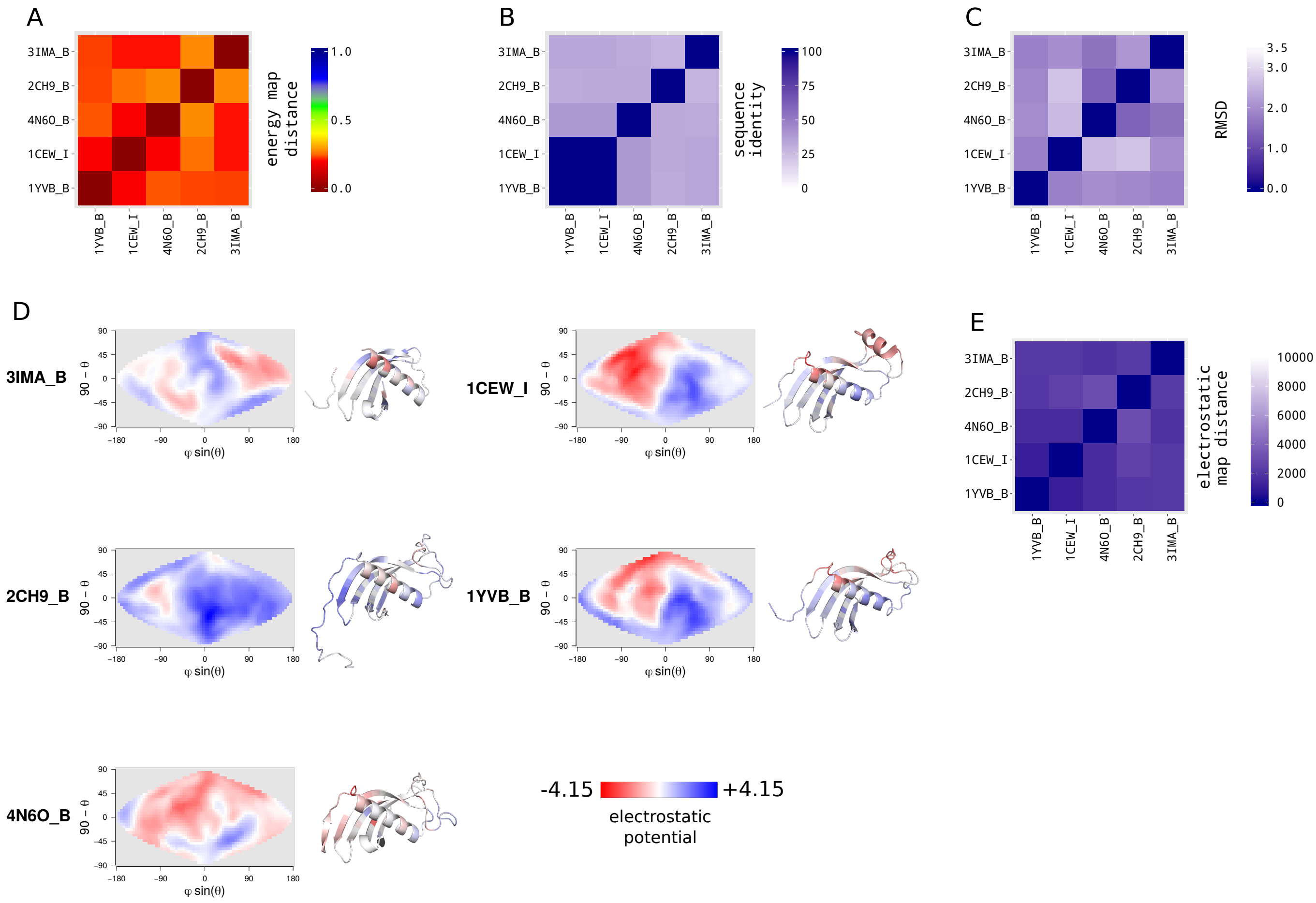


**S13 Fig. Cystatins family.** (*A*) Energy map distances matrix. It corresponds to the subsection of the ADM for the cystatins family (for the construction of the ADM, see *Materials and Methods*). Each entry *(i,j)* represents the pairwise energy map distance of the ligand pair *(i,j)* averaged over the 74 receptors of the dataset (for more details, see *Materials and Methods*). (*B*) Pairwise sequence identity matrix between all members of the family. (*C*) Pairwise root mean square deviation (RMSD) matrix between all members of the family. (*D*) Electrostatic maps and cartoon representations of the seven members of the family. An electrostatic map represents the distribution of the electrostatic potential on the surface of a protein (for more details, see Fig. S15 and *Materials and Methods*). Cartoon structures are colored according to the distribution of their electrostatic potential. (*E*) Electrostatic map distances matrix. Each entry *(i,j)* of the matrix represents the Manhattan distance between the electrostatic maps of the proteins *(i,j)*.

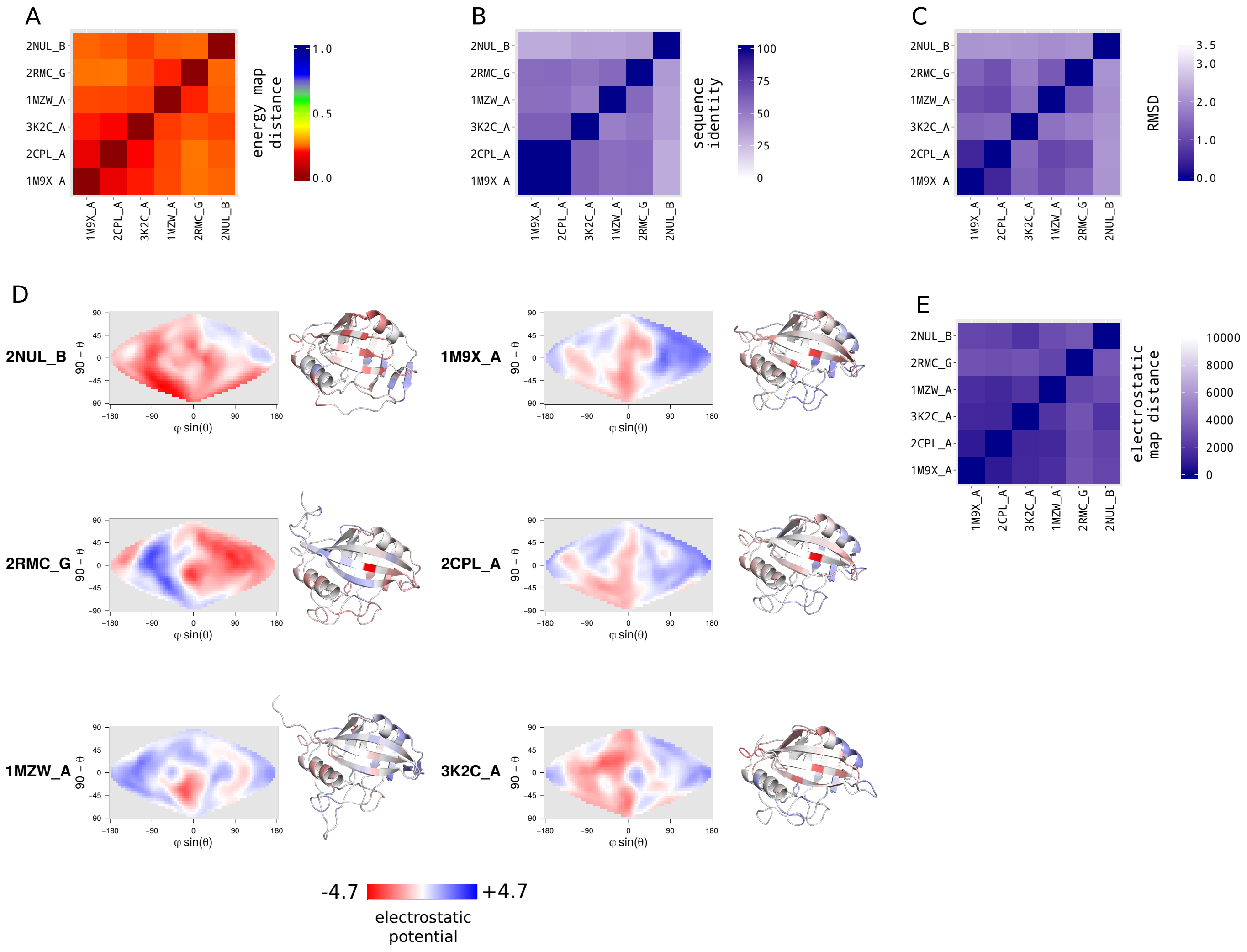


**S14 Fig. Cyclophilins family.** (*A*) Energy map distances matrix. It corresponds to the subsection of the ADM for the cyclophilins family (for the construction of the ADM, see *Materials and Methods*). Each entry *(i,j)* represents the pairwise energy map distance of the ligand pair *(i,j)* averaged over the 74 receptors of the dataset (for more details, see *Materials and Methods*). (*B*) Pairwise sequence identity matrix between all members of the family. (*C*) Pairwise root mean square deviation (RMSD) matrix between all members of the family. (*D*) Electrostatic maps and cartoon representations of the seven members of the family. An electrostatic map represents the distribution of the electrostatic potential on the surface of a protein (for more details, see Fig. S15 and *Materials and Methods*). Cartoon structures are colored according to the distribution of their electrostatic potential. (*E*) Electrostatic map distances matrix. Each entry *(i,j)* of the matrix represents the Manhattan distance between the electrostatic maps of the proteins *(i,j)*.

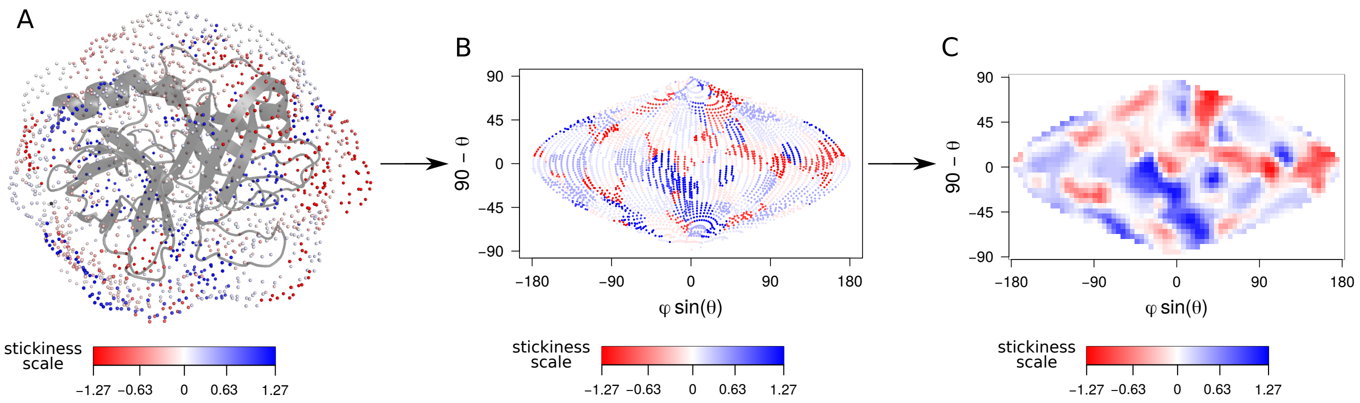


**S15 Fig. Generation of electrostatics, stickiness, hydrophobicity and circular variance (CV) maps.** Here is presented an example of generation of the stickiness map for the structure 1AVW_A. (*A*) Generation of particles with a slightly modified Shrake-Rupley algorithm [4] around the protein surface, leads to a homogenous shell of particles with a 1Å$^2$ density. Each sphere is located at 5Å from the surface of the protein. The stickiness value of the closest atom of the protein is attributed to each particle. In this example, spheres are colored according to the stickiness of the protein surface. The procedure is similar for hydrophobicity and CV. (*B*) The spherical coordinates of each sphere is represented on a 2-D map with an equal-area sinusoidal projection, following the same protocol as described in Fig. 2 and *Materials and Methods*. Each resulting dot is colored according to the same scale of (*A*). (*C*) The map is smoothed following the protocol in Fig. 2 and *Materials and Methods*. The scale is the same as in (*A*).

**S4 Table. Correlation between energy scores and stickiness, hydrophobicity and circular variance (CV)**

| | Spearman correlation | p-value |
|---|---|---|
| energy vs stickiness | -0.36 | < 2.2e-16 |
| energy vs hydrophobicity | -0.24 | < 2.2e-16 |
| energy vs CV | -0.26 | < 2.2e-16 |

The correlation is computed between each cell of the 74 energy maps of each of the 74 receptors and the corresponding cell in receptor's maps of stickiness, hydrophobicity and CV.

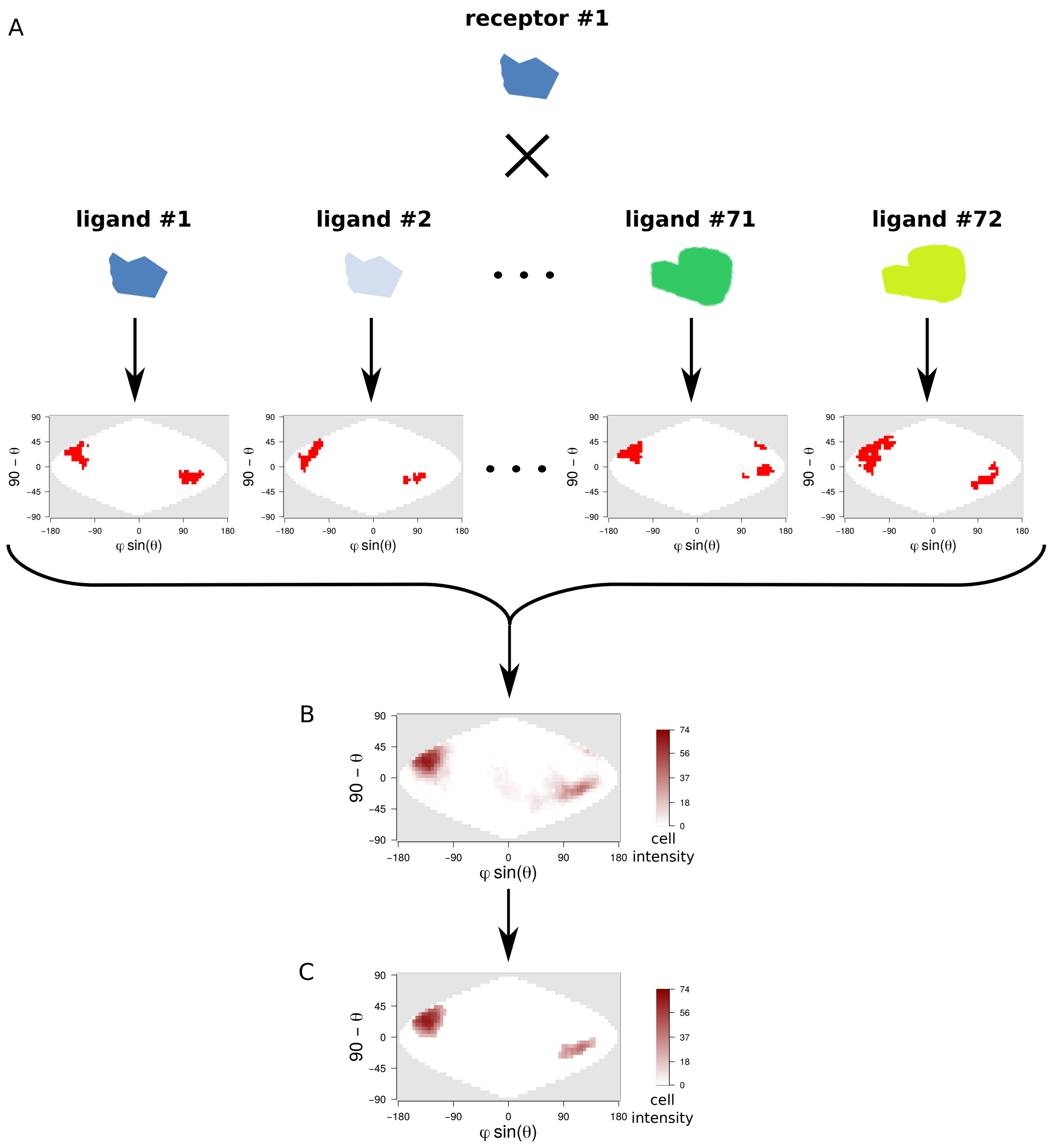


**S16 Fig. Generation of stacked maps of a receptor.** (*A*) Calculation of the 74 one-class maps (red ones in the example) of receptor #1. A value of one is associated to colored cells while zero is assigned to white cells. (*B*) Sum of the 74 one-class maps into a stacked map. Cell's intensity varies from 0 to 74 and corresponds to the number of time the cell is colored over the 74 ligands. (*C*) Filtering of the cells of low cell intensity (intensity < 17) and areas of too small size (< 4 cells) with a Dirichlet process mixture model simulation for image segmentation [5]. The procedure is repeated for each class stacked map.

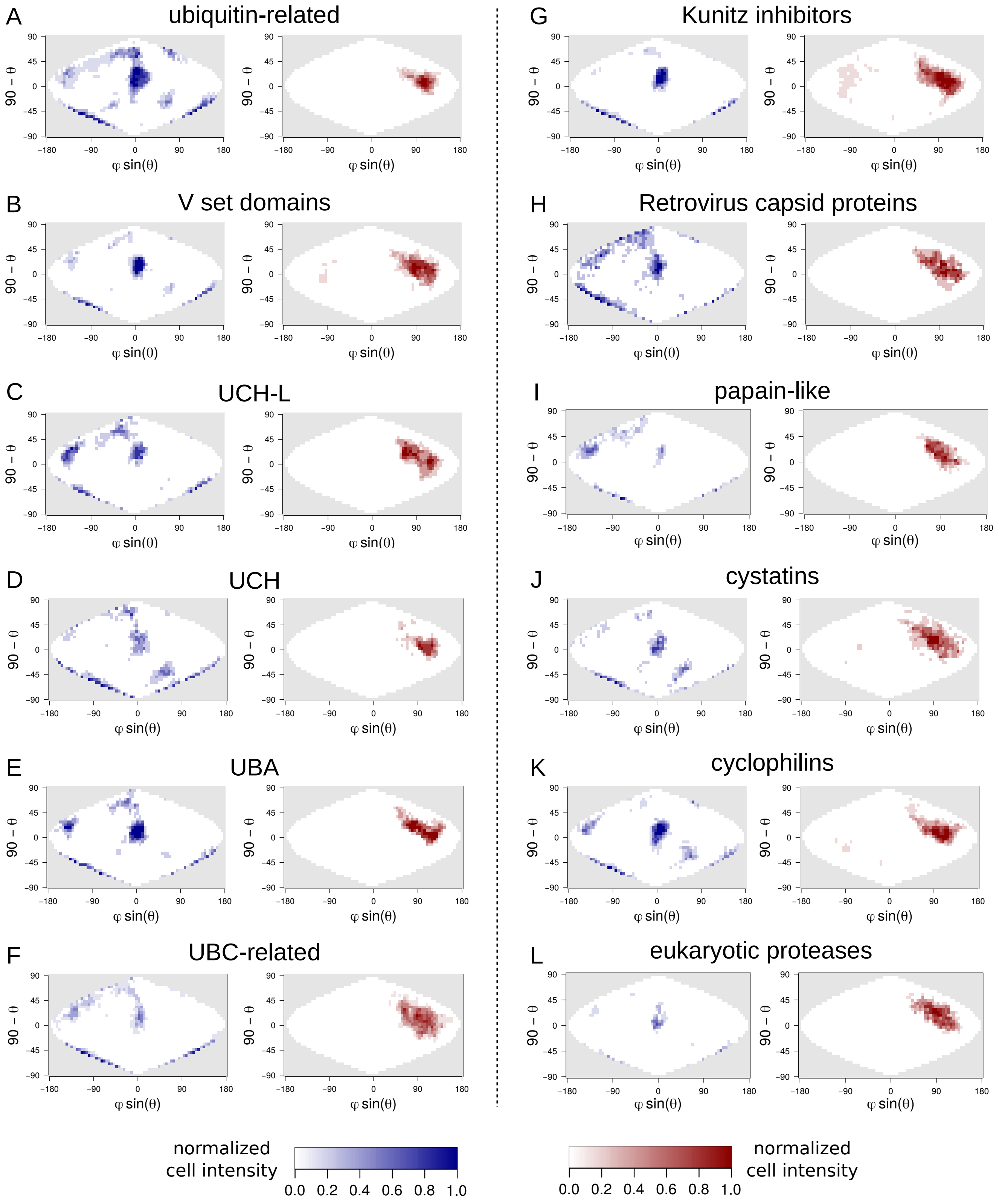


**S17 Fig. Blue and red stacked maps of 1P9D_U computed for each ligand family.** (*A-L*) We compute the one-class stacked map of each family as the sum of the one-class maps resulting from the docking of each ligand of a same family with 1P9D_U.

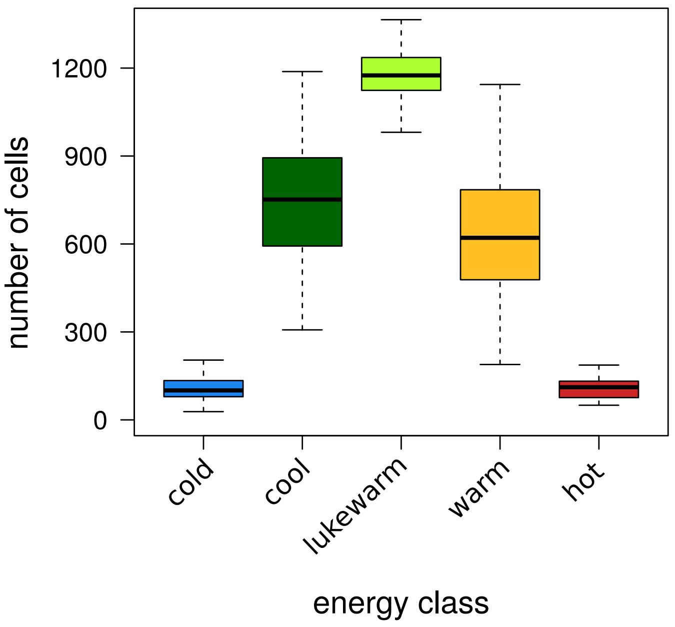


**S18 Fig. Boxplots of the size (in number of cells) of each energy class for all stacked maps.** One should notice that the sum of the sizes of the 5 energy classes is superior to 1548 cells, which is the total size of a map, because a same cell of a stacked map can be assigned to several energy classes (Fig 8).

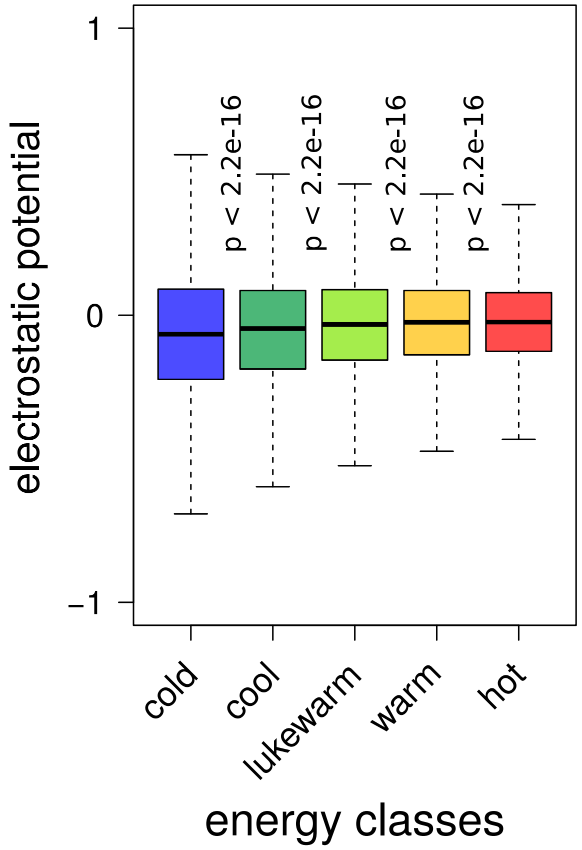


**S19 Fig. Boxplots of the electrostatic potential of the protein surfaces depending on the energy class.** The electrostatic potential is calculated for each protein following the protocol described in *Materials and Methods*. *p-values* between the variances of two "successive" energy classes were computed using the F-test.

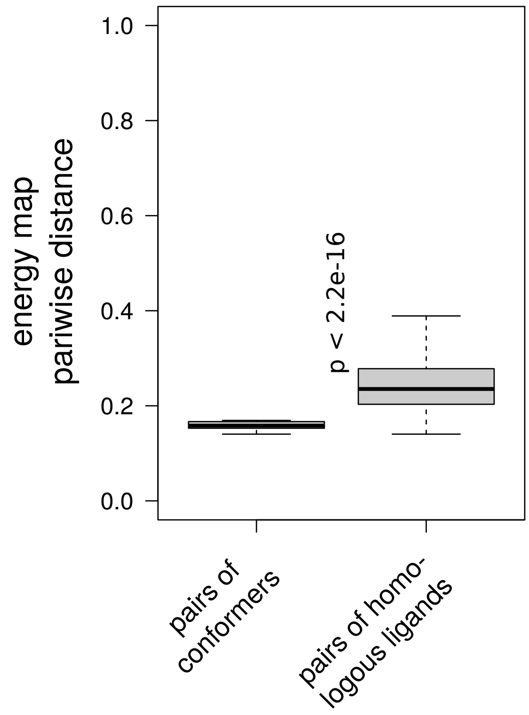


**S20 Fig. Boxplots of energy map pairwise distances between ligand pairs of conformers and pairs of homologous ligands (i.e. non-conformers pairs).** For each receptor, we computed (i) the average of energy map distances of pairs of conformers, (ii) the average of energy map distances of pairs of homologous ligands. P-values are calculated with an unilateral Wilcoxon test.

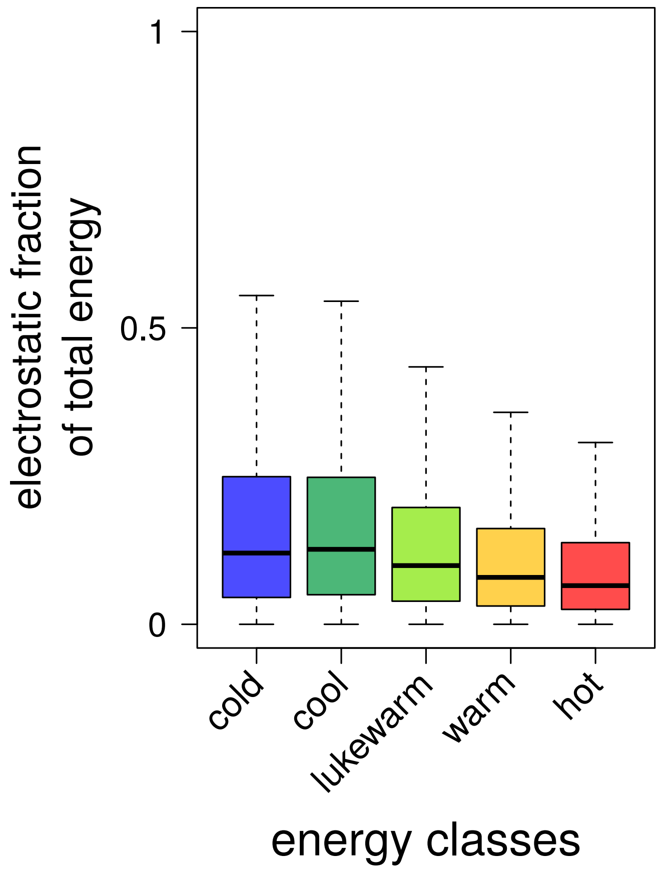


**S21 Fig. Boxplots of the fraction of the electrostatic contribution to the total energy score for each energy class.**
